# The (un)likelihood of clock-driven lateral root priming; A modeling exploration

**DOI:** 10.64898/2025.12.20.695687

**Authors:** Kirsten H. ten Tusscher

## Abstract

Lateral root formation starts with priming, predisposing subsets of pericycle cells with the potential of future lateral root formation through semi-periodic elevations in auxin (signalling). While various mechanisms have been suggested, the most broadly accepted hypothesis proposes a genetic, cell-autonomous root clock akin to the clock underlying vertebrate somitogenesis. Still, while gene expression variations were observed, so far this clock has not been proven. From a functional and evolutionary perceptive it is furthermore an open question whether lateral root priming is not more likely to arise from an emergent tissue level process similar to phyllotaxis. To help settle this debate in this study I use general knowledge of oscillator dynamics and simple models of auxin signalling to underline the unlikelihood of a root clock driving lateral root priming. I show how within a single cell oscillations are limited to a small parameter domain, with constraints intensifying due to the presence of multiple AUX/IAA and ARF types. Furthermore, I demonstrate how non-meristem based oscillations, due to a lack of memory of oscillator phase, can not drive periodic prebranch site formation.

## Introduction

Root system branching is a major determinant of a plant’s access to water and nutrients as well as its capability to anchor to its substrate. Independent of whether a plant has a taproot or fibrous root system, either the single main root or the many equivalent fibrous roots further branch through the formation of lateral roots (Figure 1A). Lateral root formation has been studied in most detail in the model plant Arabidopsis thaliana. Here the first steps in this process involve semi-periodic elevations in auxin (signalling) called priming that prepattern groups of cells to gain competence for future lateral root formation (De Smet et al., 2007; Moreno-Risueno et al., 2010) (Figure 1B). Dependent on the level of auxin elevation, a so-called stable prebranch site is formed from which in later stages lateral root initiation, primordium formation and lateral root emergence may occur. For each of these steps in lateral root formation, as well as the subsequent rate and angle of outgrowth of the emerged root, auxin signalling and environmental conditions are major determinants (Lavenus et al., 2013; Santos Teixeira & Ten Tusscher, 2019).

**Figure 1.**
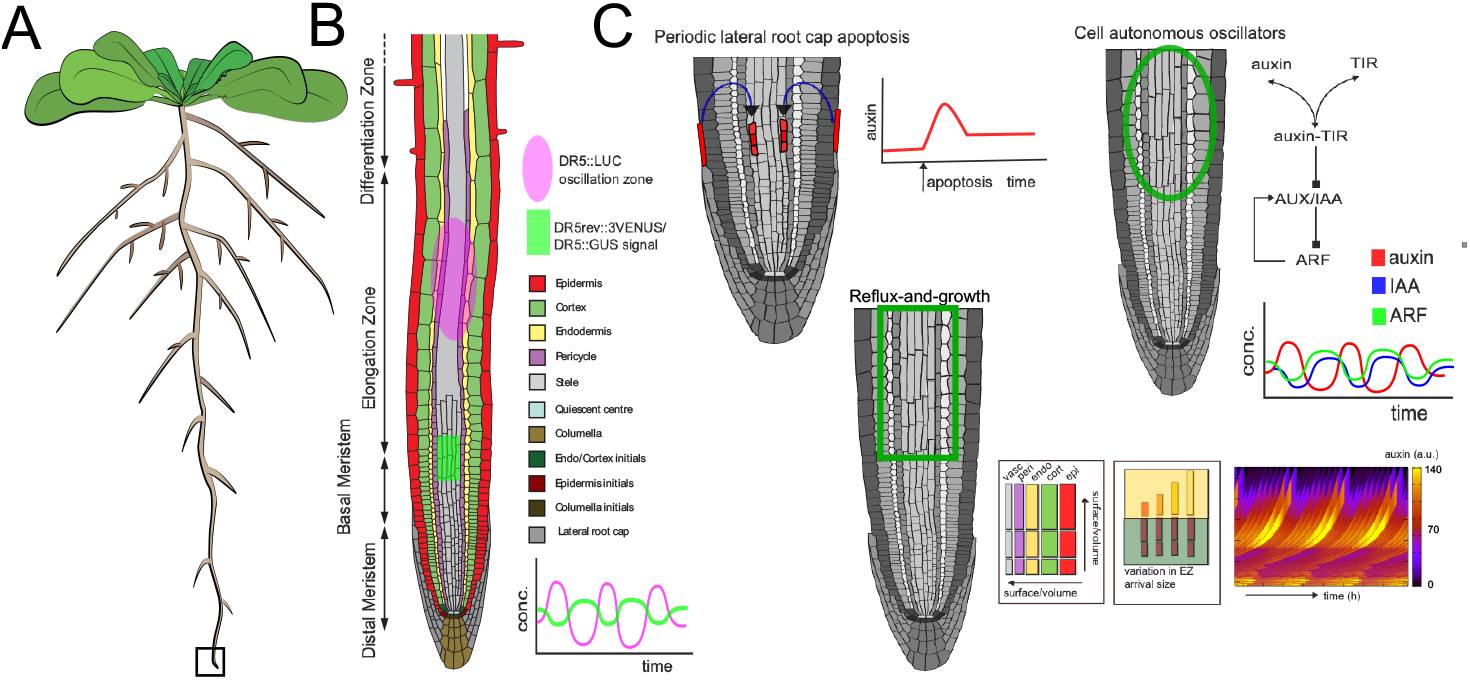
Various hypotheses for root system priming. A) A schematic of Arabidopsis thaliana rosette stage plant architecture, showing the branching of the root system through the formation of lateral roots. B) Zoom in of the boxed region in A the first signs of lateral root priming are observed just above the meristem while the oscillation zone appears to be localized more shootward. C) Three alternative hypothesis for the mechanism underlying lateral root priming, periodic root cap apoptosis, reflux-and-growth and a root clock.

The elevations in auxin (signalling) involved in priming occur above the meristem, in the elongation zone, which has been dubbed the oscillation zone (Moreno-Risueno et al., 2010). More detailed measurements indicate that these elevations occur in the vasculature just above the meristem and become transduced to the neighboring xylem pole pericycle cells from which lateral root formation takes place higher up in the elongation zone (De Smet et al., 2007; De Gernier et al., 2025) (Figure 1B). Experimental data also indicate that auxin transport and availability (De Smet et al., 2007; Xuan et al., 2015a) play a major role in lateral root priming amplitude, while the periodicity of the process strongly correlates with the process of shedding of uppermost lateral root cap cells (Xuan et al., 2016). Based on these findings models have been proposed explaining lateral root priming from root cap apoptosing cells transmitting their auxin into the main root (Xuan et al., 2016), feedback interactions between auxin concentration and auxin transporter levels (Mironova et al., 2010), or the interplay between root tip auxin reflux and stem cell driven growth (van den Berg et al., 2021) (Figure 1C). In these studies, priming arises as an emergent property from a tissue level interplay between auxin transport and growth or regulatory processes, and individual cells do not undergo oscillatory dynamics. Notably, in phyllotaxis auxin maxima have been shown to arise from the interplay between growth and auxin transport (Jönsson et al., 2006; Smith et al., 2006).

Other experimental data have shown that concurrent with the elevations in auxin signalling, a large number of genes show in or out of phase changes in expression (Moreno-Risueno et al., 2010). Involved genes drive processes such as vesicle trafficking, cell wall pectin esterification (Wachsman et al., 2020) and retinal binding proteins (Dickinson et al., 2021) that affect cell length, oscillation amplitude and lateral root formation success. The occurrence of periodic changes in gene expression led to the proposal of a so-called root clock, analogous to the well-established somitogenesis clock driving vertebrate axial patterning (Figure 1C). Such a clock would entail a non-linear, delayed negative feedback mechanism driving cell autonomous oscillations in gene expression (Novák & Tyson, 2008). Through combining such a clock with a so-called morphogen wavefront that stops oscillations while memorizing phase, these temporal oscillations become translated into a spatially periodic pattern (for the original model see (Cooke & Zeeman, 1976), for a detailed multi-scale model see (Hester et al., 2011), for recent models see review (McDaniel et al., 2024)). For somitogenesis, the molecular players underlying both clock and wavefront have been identified in considerable detail (see e.g. Aulehla et al., 2003; Dubrulle et al., 2001; Palmeirim et al., 1997). For lateral root priming, a series of modeling studies have proposed experimentally observed negative feedbacks in auxin signalling to drive gene expression oscillations. These models involve negative feedback arising from either auxin mediated freeing of ARFs inducing the expression of the ARF repressive AUX/IAA proteins (Middleton et al., 2010; Muraro et al., 2011, Muraro et al., 2013), or ARF mediated induction of auxin degredation (Mellor et al., 2016), and delays arising from slow ARF dimerisation dynamics (Middleton et al., 2010).

The semantics and citations in literature indicate that experimental plant scientists have embraced the concept of a cell autonomous root clock, and largely ignore the potential for a more tissue level emergent mechanism (see eg Bustillo-Avendaño et al., 2022; Dickinson et al., 2021; Kircher & Schopfer, 2023; Wachsman et al., 2020) with a notable exception being (Reyes-Hernández & Maizel, 2023). This broad acceptance can be understood from the fact that other clocks, such as the cell cycle and circadian clock are shared across kingdoms. Additionally, some striking similarities exist between the antagonistic FGF/Wnt and RA gradients along the vertebrate body axis and the the auxin/PLT and cytokinin signalling gradients in the root tip (K. Ten Tusscher, 2020). Still, an important question is whether similar mechanisms should be expected to be evolutionary selected for in animal segmentation and root system branching. In animals, symmetry, scaling and precision are of key importance for mobility and fitness.In plants, instead adaptability to environmental condition is essential (K. Ten Tusscher, 2020). Additionally, there are also striking differences between the two processes. Somitogenesis entails oscillations in the growth zone and individual cells undergo multiple rounds of oscillations, while lateral root priming seemingly involves only half a round of an oscillation in the elongation zone where cells no longer divide (Reyes-Hernández & Maizel, 2023; Traas & Vernoux, 2010).

Importantly, none of the modeling studies investigating the potential for a root clock have shown auxin (signalling) oscillations to arise specifically in the elongation zone from the modeled auxin signalling network and local parameter conditions. Instead, the models by (Muraro et al., 2011, 2013) suggest that auxin promotes while cytokin inhibits oscillations, implying that oscillations would occur in the meristem and terminate in the early elongation zone (Dello Ioio et al., 2007; Salvi et al., 2020), contrasting with experimental observations. Indeed, in the only modeling study simulating oscillations restricted to the elongation zone, instead of oscillations endogenously emerging from the modeled regulatory interactions, oscillatory mRNA dynamics were locally imposed (Perianez-Rodriguez et al., 2021). While absence of proof can never serve as proof of absence, we can use mathematical modeling to investigate the (un)likelihood of a cell-autonomous root clock mechanism operating from the so-called oscillation zone region to generate the periodic priming dynamics observed *in planta*. Using simple models of auxin signalling, I demonstrate how in single cells oscillations occur only in a limited parameter domain, that is further restricted by the interwinement of multiple AUX/IAA and ARF species into a single auxin signalling network and/or the addition of PIN mediated auxin export. Moving from cell to tissue level, I show that oscillations that start in the elongation zone can not drive periodic prebranch site formation due to a lack of global oscillator phase memory. While periodic inputs such as periodic root cap apoptosis may introduce phase shifting, this would take several oscillation cycles to unfold, and the stimulus rather than the oscillator would be setting the pace. I conclude that an autonomous root clock operating from the oscillation zone is highly unlikely to underly lateral root priming.

## Results

### A primer on clocks, limit cycles and oscillations

For oscillations to emerge in a biological system a so-called **negative feedback loop** is essential (Lewis, 2003; Novák & Tyson, 2008). One of the simplest examples is shown in Figure 2A, where a gene is translated into a mRNA that encodes a protein that acts as a repressive transcription factor for the expression of its own gene. Negative feedback in itself is insufficient to give rise to oscillations, instead the default behavior is **homeostasis** (Lewis, 2003; Novák & Tyson, 2008), (Fig 2B).

**Figure 2.**
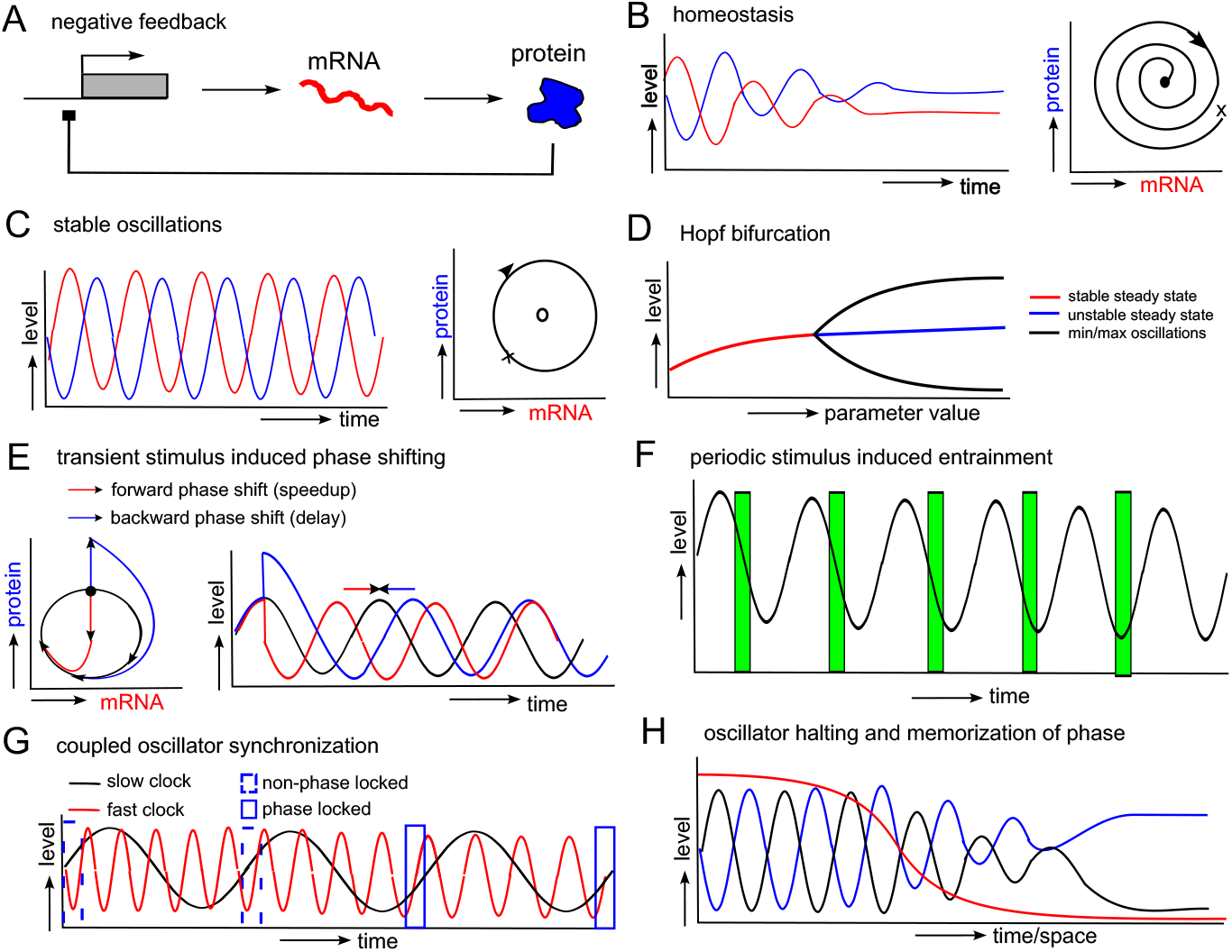
Homeostasis versus sustained oscillations, and oscillator modulation. A) A simple genetic negative feedback system in which the protein encoded by a gene negatively regulates its own expression. B) Example dynamics for a situation where negative feedback leads to homeostasis, the convergence of the system to a stable equilibrium, as a function of time (left) and in a 2D phase space (right). C) Example dynamics for a situation where negative feedback leads to stable oscillations, as a function of time (left) and in a 2D phase space (right). D) Bifurcation diagram showing how equilibrium and oscillation presence and stability change as a function of parameter value. Stable oscillations arise from a Hopf-bifurcation, involving a destabilisation of a previously stable equilibrium. E) A single stimulus can shift the phase of an oscillator forward (red) or backwards (blue), causing the system to be ahead or behind of the state it would have been in without the stimulus. F) Repeated application of a stimulus (timing indicated by the green bars) results in the eventual alignment of a particular oscillator phase (here the valley) with the stimulus. G) Coupling of oscillators with substantially different periods results in adjustments of both oscillators resulting in eventual alignment of particular phases in both oscillators, in this case the upward phase of both oscillators. H) A oscillation impacting morphogen gradient (red line) through which individual cells (black and blue lines) sequentially traverse enables memorisation of different oscillator phases in different cells. The memorisation requires an additional mechanism besides the oscillator and morphogen gradient.

Only if interactions are sufficiently **temporally delayed and input-output relations are non-linear** (Lewis, 2003; Novák & Tyson, 2008), sustained oscillations may arise (Fig 2C). Put into words, non-linearities are essential to amplify a bit too much protein in a strong decline in mRNA levels, and vice versa, causing the system to over and undershoot. Delay results in mRNA levels responding to protein levels of a while ago rather than the current instantaneous protein levels, further preventing the system to correct itself. Plotting these dynamics on a 2D mRNA versus protein plane we see that for homeostasis the system spirals inward to a stable equilibrium (Fig 2B, right), whereas for oscillations the system keeps walking over a series of connected points in this 2D plane (Fig 2C, right) like the hands that keep traversing the face of a clock. In mathematical terms this path of connected points is called a **limit cycle**. Non-linearities and delays frequently occur in biological systems. Non-linearities for example arise from cooperative binding of the transcription factor protein to its promotor, resulting in a non-linear translation of protein levels into gene expression changes. Delays arise from transcription, splicing, translation and dimerisation and transport of mRNA and protein between nucleus and cytosol (Novák & Tyson, 2008). Finally, in addition to negative feedback, delay and non-linearity, also other parameters affect oscillatory behavior. A parameter dependent transition of a system between stable and oscillatory behavior is called a **Hopf bifurcation** (Fig 2D).

### Control and coordination of clocks

While oscillators are able to autonomously maintain their cyclic behavior, they can be impacted by other factors. **Phase shifting** results from the effect of a single transient stimulus on one or more clock variables (Ermentrout & Terman, 2010; Winfree, 1980). In this case, I assume our stimulus either transiently enhances (Fig 2E left, blue) or decreases (Fig 2E left, red) protein level. In the 2D plane, we see how protein increase causes a detour while protein decrease causes a shortcut relative to the normal limit cycle (Fig 2E left), resulting in a backward and forward phase shift, respectively (Fig 2E, right). Thus, while oscillation period is unaffected, timing is. In contrast, **entrainment** results from the repeated, periodic application of a stimulus. Here, because the stimulus repeatedly pushes one or more of the systems variables in a certain direction, the system responds by aligning a particular oscillation phase with the timing of the stimulus (Figure 2F, the oscillator aligns its valley with the green stimulus). A familiar example is the entrainment of organism’s circadian clock with the light period during the day (Bell-Pedersen et al., 2005). Entrainment affects both phase and period, and if entrainment period differs more strongly from the free running period stronger stimuli are required (Ermentrout & Terman, 2010; Winfree, 1980).

Often oscillators are coupled to other oscillators, for example in multi-cellular tissue, where individual cells become **synchronized** through intercellular communication (Ermentrout & Terman, 2010; Winfree, 1980). Synchronisation becomes less trivial for oscillators with a very different frequency (Figure 2G), e.g. the cell cycle and circadian clock within a single cell. Here synchronisation implies the adjustment of frequency and phase of both oscillators, causing them to become **phase locked** to one another in a particular phase (Ermentrout & Terman, 2010; Winfree, 1980). Similar to entrainment, this requires sufficiently strong coupling, while oscillator frequencies should not deviate to far from a 1:n ratio. Finally, as for example in the case of vertebrate somitogenesis oscillations may need to terminate and be transduced into a stable spatially periodic pattern (Fig 2H). This typically involves a gradual change in a parameter controlling oscillations, like the FGF/WNT morphogen gradient in somitogenesis. While terminating oscillations can be achieved by reducing delays, or by preventing the production of the repressive protein, an additional mechanism is required for **memorizing oscillator phase** prior to or coincident with full oscillator termination (see e.g. François & Mochulska, 2024).

### Auxin signaling drives oscillations within a constrained parameter range

The root clock underlying lateral root priming is assumed to arise from oscillations in auxin signalling and/or auxin levels. Canonical auxin signalling involves the binding of an auxin molecule to a single TIR/AFB receptor molecule (Kepinski & Leyser, 2005), with the resulting auxin-TIR/AFB complex binding to an AUX/IAA protein (Kepinski & Leyser, 2004), forming an auxin-TIR/AFB-AUX/IAA complex in which the AUX/IAA protein can be ubiquitinated, tagging it for degradation (Figure 3A). Free non-ubiquitinated AUX/IAA protein can bind to Auxin Response Factors (ARFs), preventing ARFs from inducing auxin dependent gene expression (Weijers et al., 2005). Auxin thus induces auxin dependent gene expression by derepressing ARFs. ARFs are found to homodimerize, with dimers being more potent inducers of gene expression (Fontana et al., 2023; Ulmasov et al., 1999). For at least a subset of AUX/IAA genes, ARFs induce expression of these genes, resulting in a negative feedback loop (Middleton et al., 2010; Vernoux et al., 2011).

**Figure 3.**
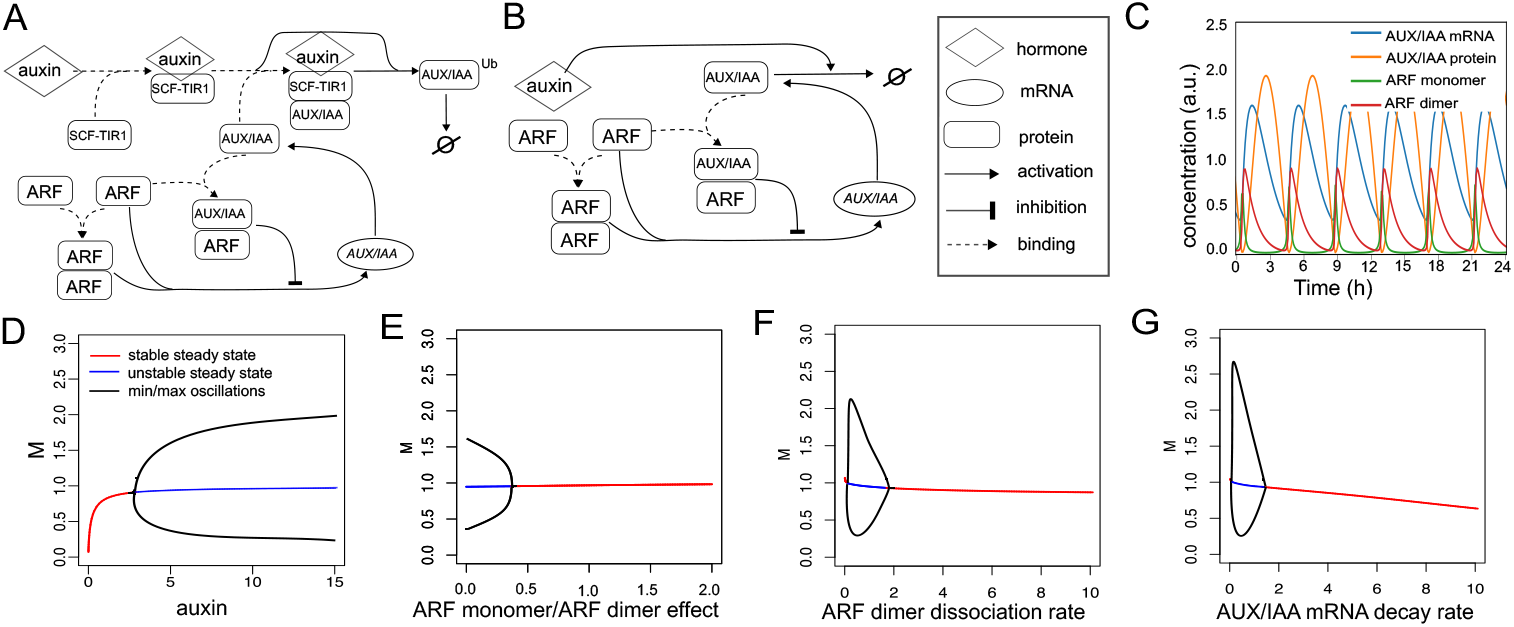
Oscillatory capacity of a single AUX/IAA single ARF system. A) Overview of the major players and interactions in plant auxin signalling as modeled in the Middleton et al. 2010 model. B) Overview of the simplified auxin signalling model in this work. C) oscillatory dynamics generated by the model shown in B for the parameter settings shown in Supplementary Table 2. D-G) Bifurcation diagrams showing dependence of oscillatory dynamics on auxin level (D), the ratio of effectiveness in inducing AUX/IAA gene expression of ARF monomers versus dimers that affects non-linearity (E), the rate of ARF dimer dissociation that affects delay duration (F) and the AUX/IAA mRNA decay rate (G). In D-G, except for the bifurcation parameter, parameters were identical to those in C.

Several similar models of the auxin signalling system have been made (Middleton et al., 2010; Vernoux et al., 2011), and here I developed a simplified version of the Middleton 2010 model (Fig 3B) (See Methods). As for the original model, for specific parameter conditions oscillatory dynamics can be obtained (Fig 3C), yet in addition to reductions in auxin levels (Fig 3D) reductions in non-linearity (the size difference in effectivenes of ARF dimers versus monomers on transcription), temporal delays in the negative feedback loop (dimer dynamics) and differences between AUX/IAA mRNA versus protein turnover readily transform oscillatory into steady state dynamics (Fig 3E-G). These results are consistent with earlier results (Vernoux et al., 2011), as well as general oscillator theory (Novák & Tyson, 2008). Thus, in a single ARF single AUX/IAA system oscillations occur in a limited parameter domain that may not trivially occur *in planta*.

### A two ARF-two AUX/IAA system further constrains oscillation capacity

Importantly, Arabidopsis contains a total of 23 ARFs and a total of 29 AUX/IAAs, and in most plant tissues a combination of several ARFs and AUX/IAAs are simultaneously present (Rademacher et al., 2011; Vernoux et al., 2011; Weijers et al., 2005). To investigate the impact on oscillatory dynamics I considered a two ARF two AUX/IAA system, assuming that the different ARFs could heterodimerize with one another, as well as with both AUX/IAAs (Fig 4A). I start from a situation in which only one ARF AUX/IAA pair is expressed (green, A), parametrised such that dynamics are well within the oscillatory regime (Supplementary Table 2). After reaching stable oscillations I allow expression of the second ARF AUX/IAA pair (red, B) parametrised to not oscillate (Supplementary Table 2). If maximum AUX/IAA and ARF protein levels of pair B are 25% or more than of pair A, oscillations were halted (Fig 4B).

**Figure 4.**
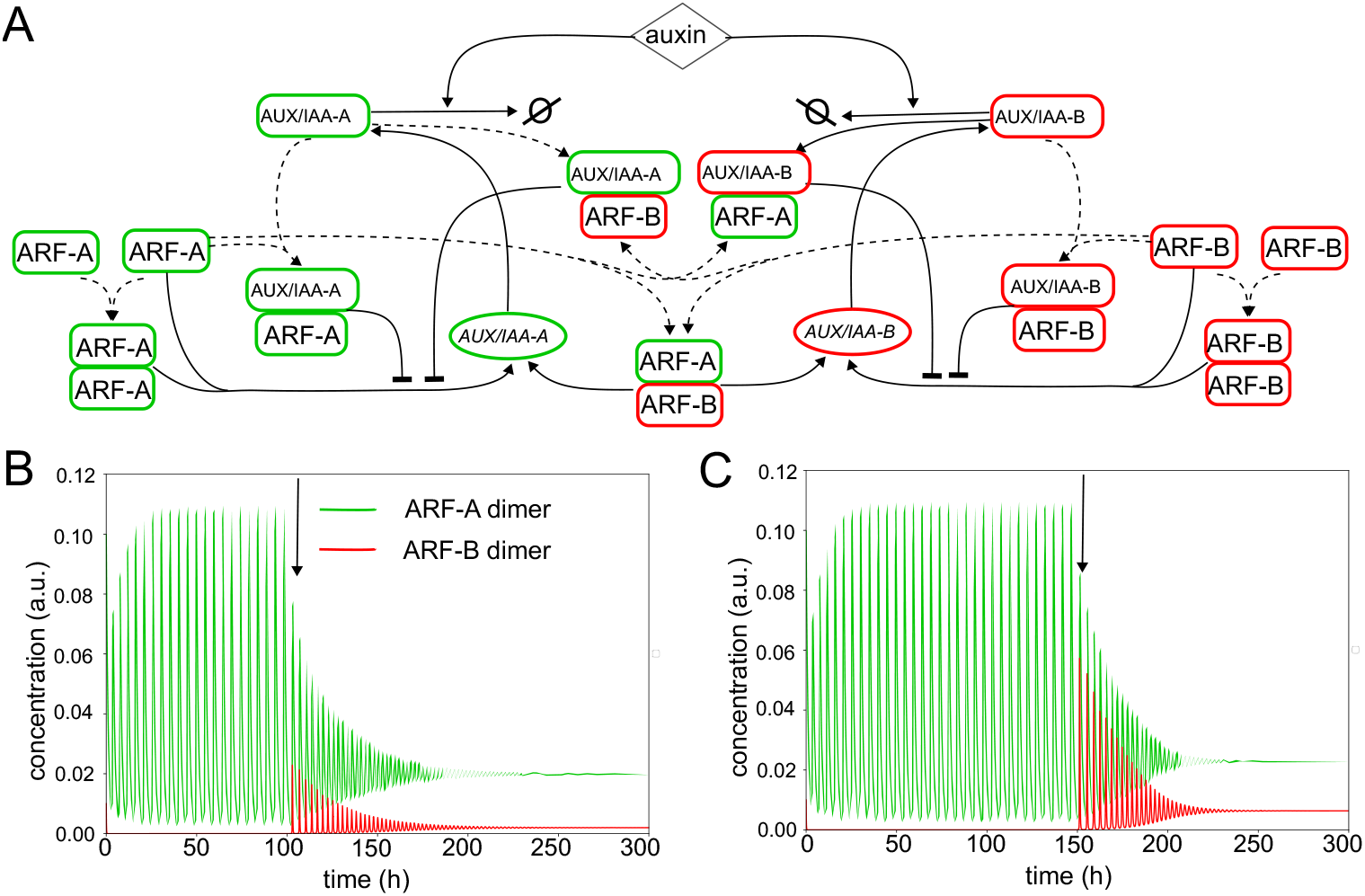
Termination of oscillations in a two AUX/IAA two ARF system. A) Architecture of a two AUX/IAA two ARF auxin signalling network in which both AUX/IAAs can dimerize with both ARFs and ARFs can heterodimerize. The “A”-pair (indicated in green) is parametrized to support oscillations (Supplementary Table 2), the “B”-pair (indicated in red) is parameterized to not support oscillations (Supplementary Table 2) B) System dynamics during an initial phase in which only the “A” pair is expressed, and a second fase (start indicated by the arrow) where also the “B” pair is expressed C) Same as in B but now with ARF-A monomers and dimers having 20-fold less affinity for AUX/IAA-B promotors than ARF-B monomers and dimers, and ARF-B monomers and dimers having 20-fold less affinity for AUX/IAA-A promotors than ARF-A monomers and dimers. ARF heterodimers have 20-fold lower affinity for both AUX/IAA promotors.

Still, *in planta* ARFs and AUX/IAAs may, in addition to e.g. turnover parameters and dimerisation rate that affect the potential of an ARF-AUX/IAA pair to display oscillatory dynamics, also differ in binding affinities and effects on target genes. Therefore I redid the above simulation, with ARF A monomers and dimers mostly affecting expression of AUX/IAA A and ARF B mostly affecting expression of AUX/IAA B through implementing a 20-fold difference in the binding affinities of ARFs to the different AUX/IAA promotors. Keeping all else identical, the ARF-AUX/IAA B pair now needed to be expressed at at least 40% of the ARF-AUX/IAA A pair to halt oscillations (Fig 4C). Thus, even with a partial separation between the two pairs, oscillations can be easily terminated through the addition of a second, numerically non-dominant non-oscillatory ARF-AUX/IAA pair.

This indicates, that *in planta*, with multiple AUX/IAAs and ARFS parameter constraints need to at least be partially met by all AUX/IAAs ARFS in the oscillation zone to support oscillations. Thus further underlines the non-trivality of meeting these conditions.

### A degradation and import based auxin signalling oscillator can only function in absence of significant auxin export

In addition to fully auxin signalling based networks displaying oscillations, alternative models have been proposed. Here oscillations arise not from the delays and non-linearity in the ARF-dimer dominated induction of AUX/IAA gene expression, but form ARF induced LAX3 and GH3 expression (Mellor et al., 2016). Negative feedback arises from the GH3 expression, which degrades auxin, while a counteracting positive feedback arises from the auxin importer LAX3 (Fig 5A). Combining negative feedback with positive feedback is a well known method to achieve oscillations (Novák & Tyson, 2008). In isolated cells, under constant external auxin conditions and specific parameters this model will result in sustained oscillations (Fig 5B, Supplementary Table 3) (see also Mellor et al., 2016). Note that here oscillations in auxin signalling necessarily involve oscillations in auxin concentration (Fig 5B). Importantly, pericycle and protoxylem cells, where priming is assumed to occur, express polarly localized auxin exporting PIN proteins. Even when assuming PIN expression levels to be 10% of those of LAX levels, and have 70-fold lower auxin conductance, incorporation of PINs abolishes oscillations (Fig 5C).

**Figure 5.**
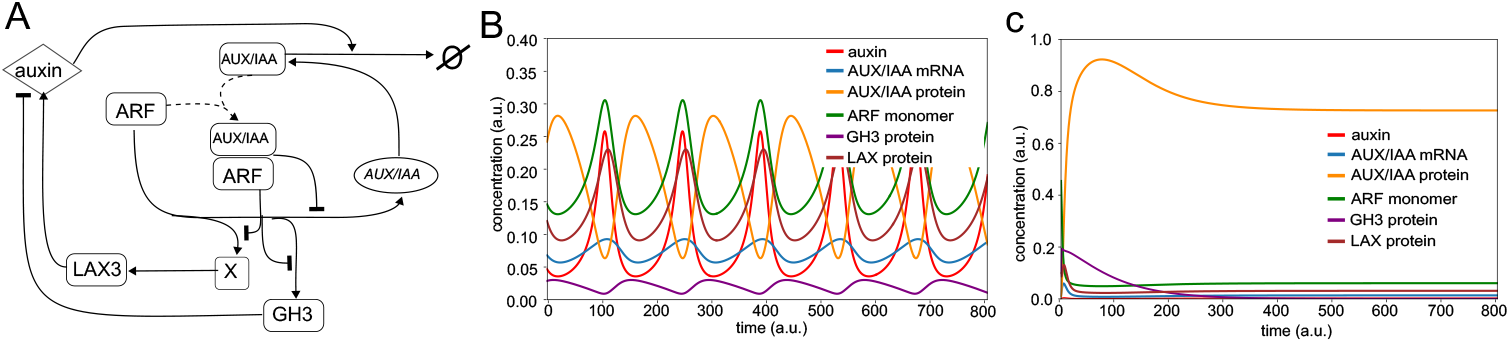
Termination of oscillations in a degradation-import oscillator by PIN efflux. A) Interaction network of auxin signalling, GH3 mediated degradation and LAX3 mediated import as modeled in Mellor et al., (2016). B) Oscillatory dynamics resulting from the parameter settings described in Table 3. C) Non-oscillatory dynamics resulting from the same parameter settings but adding a small PIN mediated auxin efflux term.

### A model for tissue level evaluation of the root clock hypothesis

To further investigate constraints of an auxin-signalling root clock driving lateral root priming in a root tissue context, I from now on simply assume a single ARF-AUX/IAA pair that -at least for a certain region in the root-is able to generate stable oscillations. I simulate a 1D tissue representing a single protoxylem and/or pericycle pole (Fig 6A), in which each cell contains the auxin signalling network (Fig 6C), and in which cells in the meristematic region divide, in the transition zone slowly grow and in the elongation zone rapidly expand (Fig 6B). In the differentiation zone cell expansion has terminated and cells maintain their size. By varying parameter settings along the 1D tissue strand one can influence where in the tissue oscillations occur (Fig 6D). In addition to the auxin signalling network, a simplified memorization mechanism is implemented that beyond a certain point in the tissue and within a limited spatial window memorizes the state of the auxin signalling network to enables transformation of temporal oscillations into spatial stripes, i.e. prebranch sites (PBS). Cell growth, division and expansion rates, meristem size and differentiation rates were fitted to reproduce recent detailed growth tracking data from Goh et al. (2023) (compare Fig 6E-F to Fig 4A in Goh et al., 2023) (see Methods, Supplementary Table 4). In the model a larger region of the root is simulated than tracked in the experiment (2.5mm versus 600microm), allowing one to further track cell size variations, growth dynamics and PBS formation into the differentiation zone, for which typically downward oriented kymographs are used (Fig 6G).

**Figure 6.**
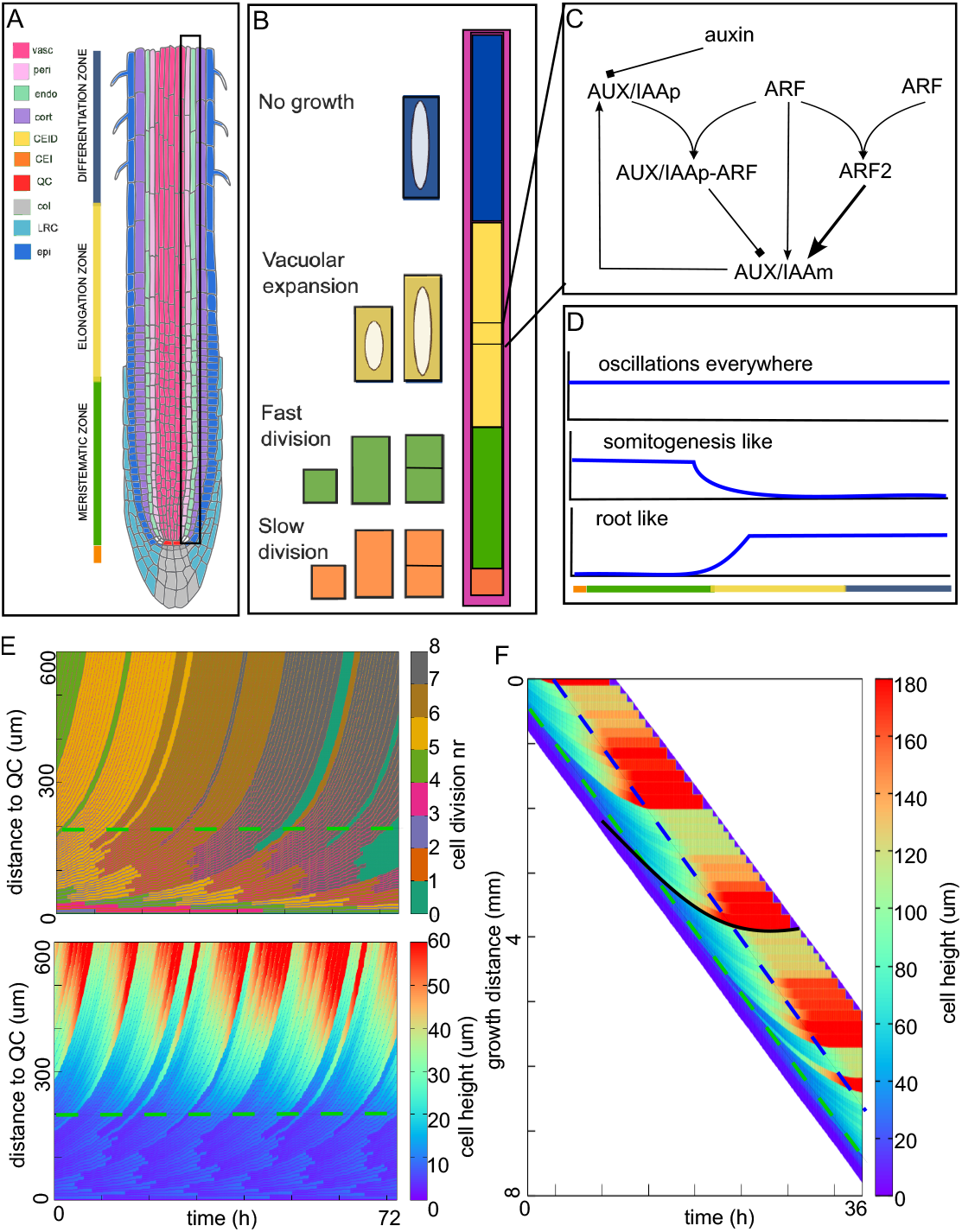
1D growing tissue model with oscillatory network. A) Overview of Arabidopsis root tip anatomy with the different cell types organized largely in cylindrical fashion and the different longitudinally organized developmental zones. Pericycle and nearby protoxylem cell files are indicated by the boxed in area. B) Layout of the 1D tissue model with developmental zones and corresponding growth dynamics. C) Simplified representation of the auxin signalling network incorporated in all individual cells. D) Possible parameter gradients (blue) governing where oscillations can occur (high level) and cannot occur (low level). E) Cell dynamics kymograph where cells are colored based on the number of cell divisions they underwent (top) or their cell height (bottom) G) Cell size dynamics kymograph from the same model simulation as in F but now showing a larger domain of the simulated root and with the root tip following actual growth trajectory instead of its position aligned to the horizontal axis. The black line indicates the trajectory of an individual cell. Dashed green lines indicate the top boundary of the meristem, dashed blue lines indicate the top boundary of the elongation zone.

### MZ localized oscillations can drive periodic PBS formation

As a first step, parameter conditions are used in which oscillatory dynamics are only supported in the meristematic growth zone. Note that this situation resembles the situation in vertebrate somitogenesis and arthropod clock-driven segmentation where oscillations occur in the posterior growth zone and terminate anteriorly. One can see a clear temporal activity pattern arises in the meristem, indicating that all meristematic cells are oscillating in phase (Fig 7A). Upon entering the elongation zone, these oscillations dampen and terminate (Fig 7A). After adding a mechanism memorizing auxin signalling state at the start of the elongation zone, this transient temporal oscillatory pattern becomes transduced into a spatially periodic pattern in the rest of the root (Fig 7B-C). To further illustrate this, Figures 7D and 7E show dynamics of 2 individual cells, their trajectories indicated in Figures 7A and 7B, as a function of either time or distance. The figures compare non-memorized (dashed lines) with memorized dynamics (solid lines), again showing how as different cells enter the elongation zone sequentially and hence at different phases (Figure 7D), different memorized auxin signalling states ensue. This enables periodic patterning of PBS, as observed in experiments.

**Figure 7.**
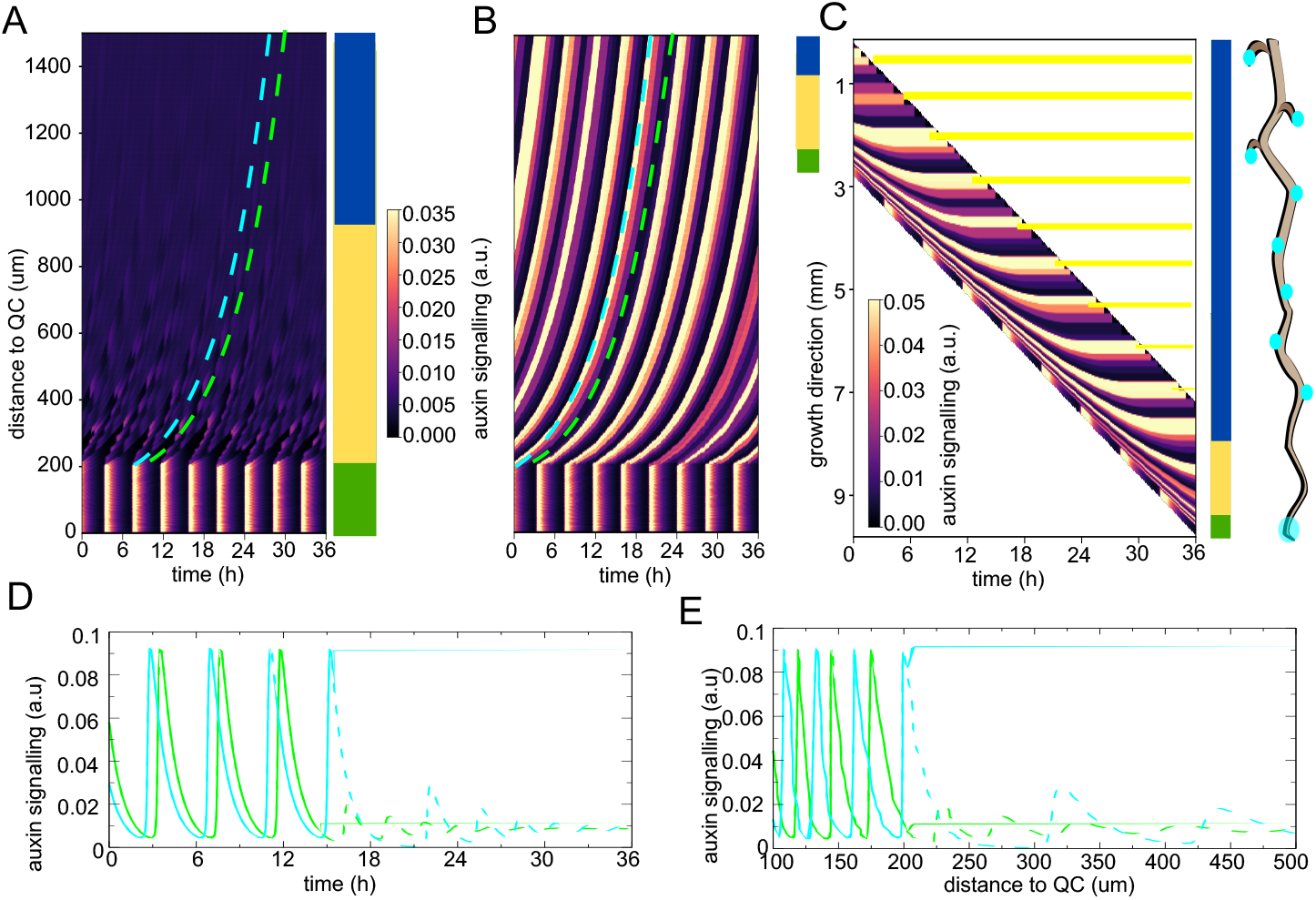
MZ driven oscillations periodically patterning PBS. A) Kymograph depicting auxin signalling dynamics (ARF dimer level) for a situation where the parameter controlling the total amount of ARF in the system is lowered from MZ to EZ such that oscillatory dynamics are only supported in the MZ. For ease of orientation the root zones to which the different distances from the QC correspond are depicted. B) Kymograph depicting auxin signalling dynamics (MZ) and subsequent memorisation (from start of EZ upwards). C Same kymograph as in B, but now for root tip displacing downward during growth. Again for orientation root zones are indicated. Furthermore, yellow lines indicate how high auxin peaks would continue if the simulation domain would keep growing like a real root instead of having a constant finite size, with the root on the right indicating how this would result in PBS and lateral root primordia patterning. D) Auxin signalling dynamics in two separate cells (cyan and green lines), in absence (dashed lines) and presence (continuous lines) of memorisation dynamics, as a function of time. Trajectories of these cells are indicated in A and B. E) Same dynamics as in D but now as a function of distance from QC.

Importantly, *in planta*, oscillatory activity related to priming has only been observed either just above the meristem (De Gernier et al., 2025; De Smet et al., 2007), or higher up in the elongation zone (De Gernier et al., 2025; Moreno-Risueno et al., 2010; Xuan et al., 2015a, 2016) depending on the reporter used, but not inside the meristem itself.

### Above MZ driven oscillations can not periodically pattern PBS

Therefore, I next implemented “root-like” conditions, where oscillatory dynamics are only supported above the meristem. Consistent with this oscillations start in the elongation zone, with cells that at different time points reside at the same physical distance from the QC displaying the same oscillation phase, resulting in a horizontally banded spatial pattern in the (unmemorized) kymograph (Figure 8A). Observed small variations in the horizontal direction arise from stochasticity in cell growth and division causing minor timing differences in when a cell enters the elongation zone. This dramatically different behavior arises from the fact that while individual cells still oscillate, they now all enter the elongation zone from the same non-oscillatory condition, starting oscillations in the same phase (Fig 8D and 9E). Applying memorization mechanism approximately halfway the elongation zone, either all cells end in a high (Figure 8B) or low auxin signalling state (not shown), depending on the exact positioning of the memorisation (bottom versus top arrow in Fig 8A).

**Figure 8.**
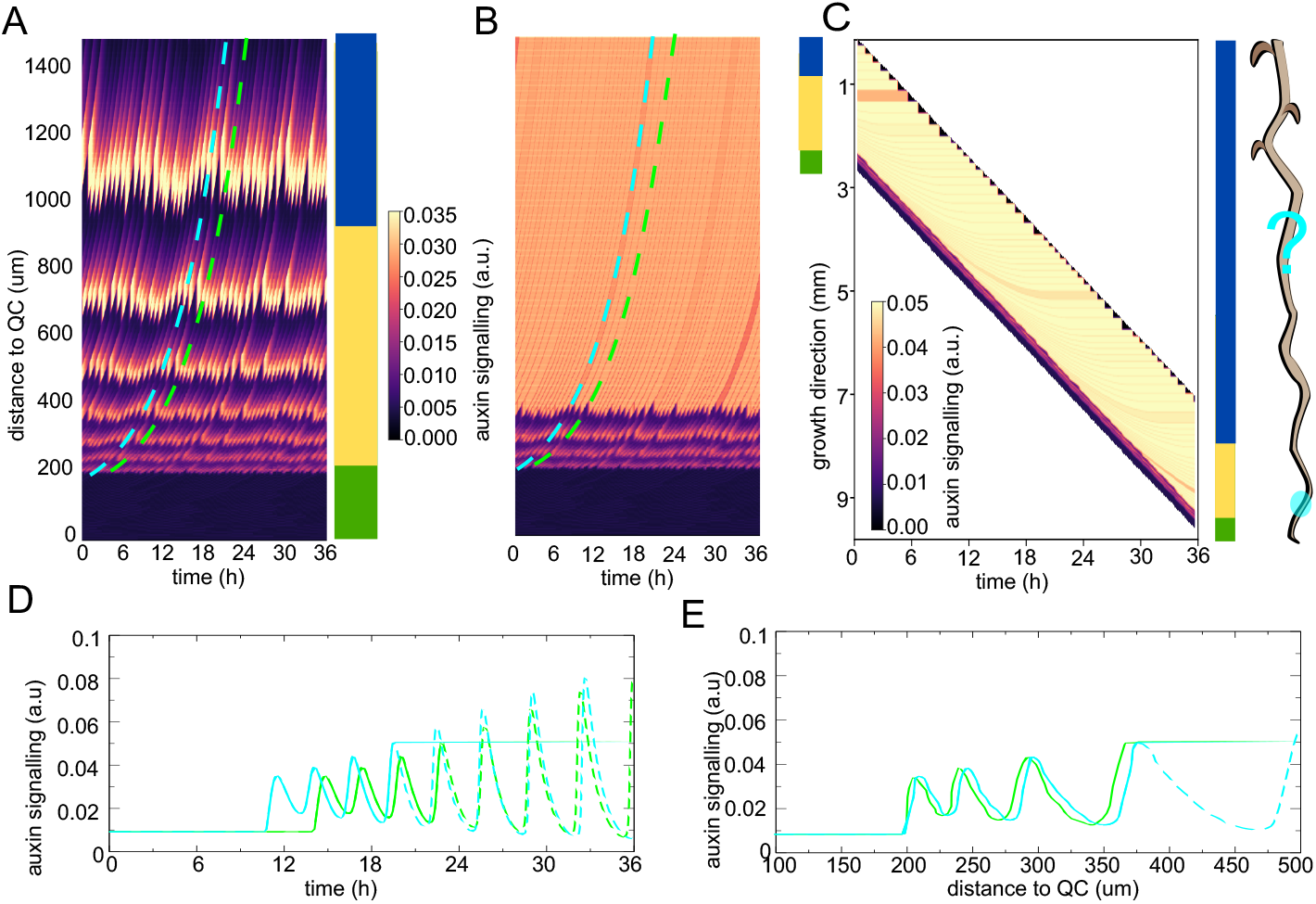
EZ localised oscillations result in spatial synchronisation. A) Kymograph depicting auxin signalling dynamics (ARF dimer level) for a situation where the parameter controlling the total amount of ARF in the system is increased from MZ to EZ such that oscillatory dynamics are only supported in the EZ and above. B) Kymograph depicting auxin signalling dynamics (MZ) and subsequent memorisation (from halfway EZ upwards, bottom arrow in A). C) Same kymograph as in B, but now for root tip displacing downward during growth. D) Auxin signalling dynamics in two separate cells (cyan and green lines), in absence (dashed lines) and presence (continuous lines) of memorisation dynamics, as a function of time. E) Same dynamics as in D but now as function of distance from QC.

These results reveal a fundamental problem with the applicability of the somitogenesis clock mechanism to plant roots. In somitogenesis and our simulation with meristem localised oscillations, oscillations occur in the stem cell niche and division zone. Daughter cells inherit their oscillatory state from their mother cell, with a reservoir of cells remaining in the stem cell niche that maintain the oscillation history and pass on the current phase of the cycle. As a consequence, there is tissue level time keeping and cells sequentially entering the EZ do so with different phases (Suppl Fig 1A). In contrast, if as observed *in planta* oscillations start only in the EZ zone, no tissue level time keeping exists and instead all cells start oscillating from the same phase as if pressing their stopwatch when passing the EZ start mark (Suppl Fig 2A). These results follow from general oscillator dynamics, independent of oscillator details, cell-cell coupling, gradients or noise.

### Global oscillator input can drive periodic PBS formation

The above may explain why in the Perianez-Rodriguez et al. (2021) study, which simulated oscillations specifically in the early elongation zone, oscillatory mRNA dynamics needed to be superimposed in this zone. To validate this hypothesis, I took a similar approach, using non-oscillatory parameter conditions (Supplementary Table 2) and replacing ARF mediated IAA transcription with superimposed oscillatory transcription (equation 20b Methods). Our model output indicates that periodic elevations in auxin signalling in the early elongation zone now could indeed be obtained (Fig 9A). Without a memorization mechanism, oscillations continue and individual cells undergo multiple rounds of oscillations (Fig 9A,D, E), while adding a memorization mechanism allows cells arriving at different phases of the global oscillator to generate a spatially periodic pattern (Fig 9B-E).

**Figure 9.**
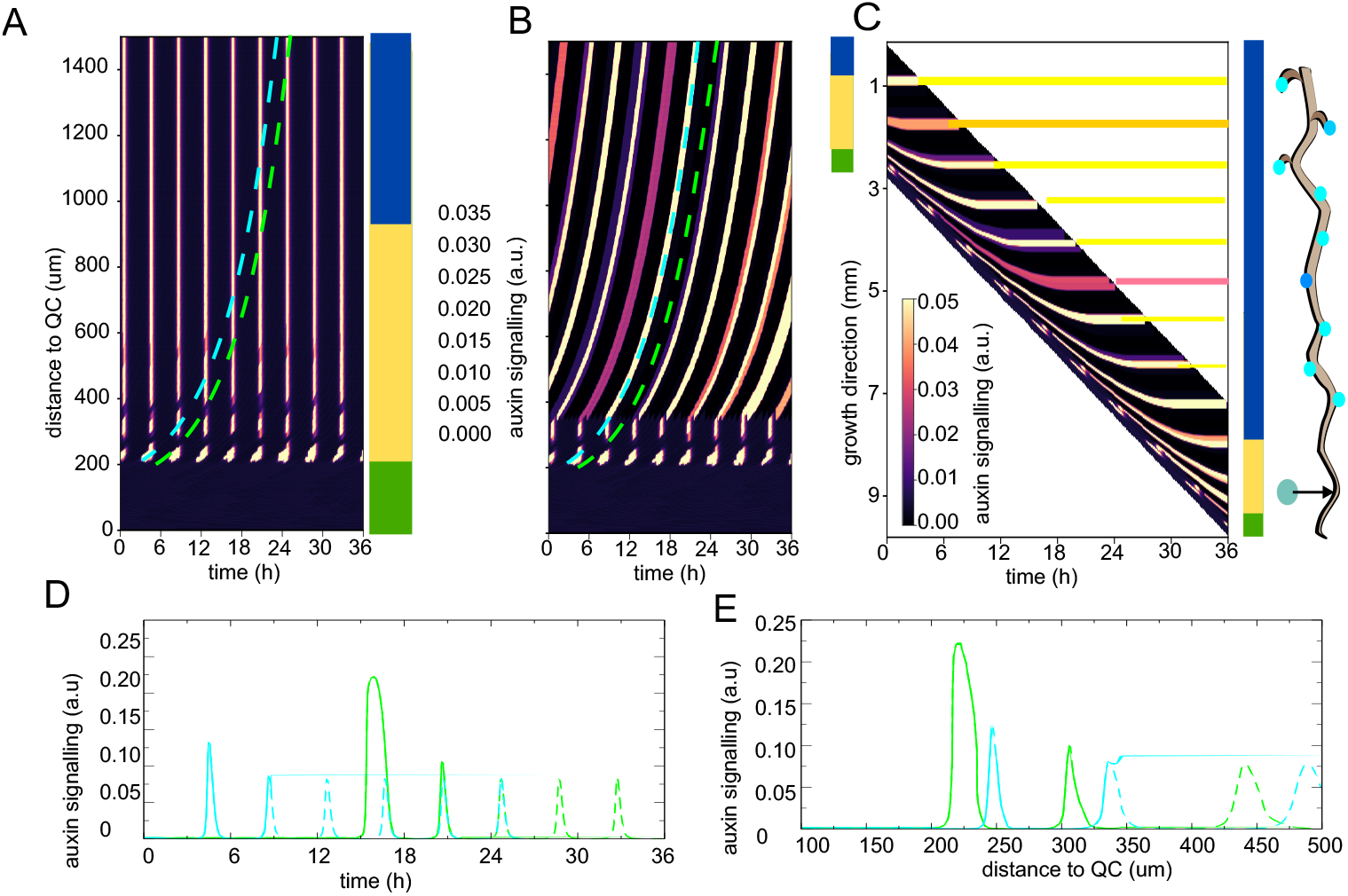
Superimposed AUX/IAA oscillations in EZ drive periodic PBS formation. A) Kymograph depicting auxin signalling dynamics (ARF dimer level) for a situation where AUX/IAA expression dynamics are superimposed to be sinusoidsal, and total amount of ARF does not support autonomous oscillations. In the root on the side illustrating the eventual result in PBS the oscillatorary zone is put aside the root, to indicate it’s superimposed rather than endogenous nature. B) Kymograph depicting auxin signalling dynamics (MZ, early EZ) and subsequent memorisation (from halfway EZ upwards. C) Same kymograph as in B, but now for root tip displacing downward during growth. D) Auxin signalling dynamics in two separate cells (cyan and green lines), in absence (dashed lines) and presence (continuous lines) of memorisation dynamics, as a function of time. E) Same dynamics as in D but now as function of distance from QC.

Importantly, by superimposing IAA mRNA oscillations, I effectively globally dictated oscillations rather than allowing them to naturally emerge from the interactions in the auxin signallling network. Note that it would be entirely unclear where this global oscillation phase information would be stored, given that individual cells do not persistently reside in the EZ. Additionally, by superimposing IAA mRNA oscillations one is not simply providing a periodic input signal to the root clock but rather magically driving the expression of one of its genes.

### Limited rescue of periodic patterning from periodic sources

Next, I therefore investigated to what extent periodic input could enable periodic variations in the elongation zone. Since cells only start oscillating in the EZ and it is the first peak/valley that is being transduced to a PBS/no PBS, time is insufficient for entrainment (Fig 2F,G) and periodic signals can only provide a single phase shifting input (Fig 2E). Therefore, I considered periodic processes that operate at periods close to the priming period near the start of the EZ. Candidate mechanisms are root cap apoptosis resulting in export of auxin from the root cap to the rest of the root (De Gernier et al., 2025; Xuan et al., 2015b), periodic changes in elongation zone cell sizes and hence auxin uptake potential (van den Berg et al., 2021), and possibly cell cycle state given the recent discovery of faster cell cycles in the top part of the meristem (Echevarria et al., 2025). Additionally, periodic phloem unloading could be a candidate mechanism, but so far this has been reported to occur at far higher frequences (Ross-Elliott et al., 2017). I therefore tested whether a periodic variation in auxin input in the bottom 20% of the elongation zone, spanning approximately 3-4 early elongation zone cells, could drive variations in oscillator phase in the elongation zone (equation 41 Methods).

Figure 10A shows a kymograph when applying an auxin stimulus at a frequency close to the frequency of the auxin signalling driven oscillations (4.2h period). Comparing along the horizontal direction whether changes in activity occur, we see that at positions just above the meristem hardly differences occur, while higher up groups of cells become more out of phase. Tracking oscillation dynamics in individual cells (Fig 10B), we indeed see that while the auxin stimulus has an instantaneous effect on auxin signalling levels (grey area in Fig 10B), phase shifting takes several cycles to unfold. While I can not exlude the existence of stimulus parameters for which substantial phase shifting occurs instantaneously, this delayed unfolding of a phase shift is a well known property of non-linear oscillators (Kuramoto, 2003; Winfree, 1980), making this highly unlikely. As a consequence, if memorisation of auxin signalling state occurs close to the meristem (bottom arrow in Fig 10A), memorised states show limited differences between cells (Fig 10C-D) making it hard to distinghuish prebranch from non-prebranch sites. If memorisation occurs later (middle and top arrows in Fig 10A), once several cycles of oscillations have occurred and phase differences have developed makes this distinction clearer (Figure 10E-F), and PBS numbers align with the number of stimuli fitting within the time window (approximately 8 for a 4.2h period in 36h). Note that depending on the exact memorisation position, different cells will memorize a high auxin signalling state (compare Fig 10E and F), and patterning is irregular in both spacing and amplitude.

**Figure 10.**
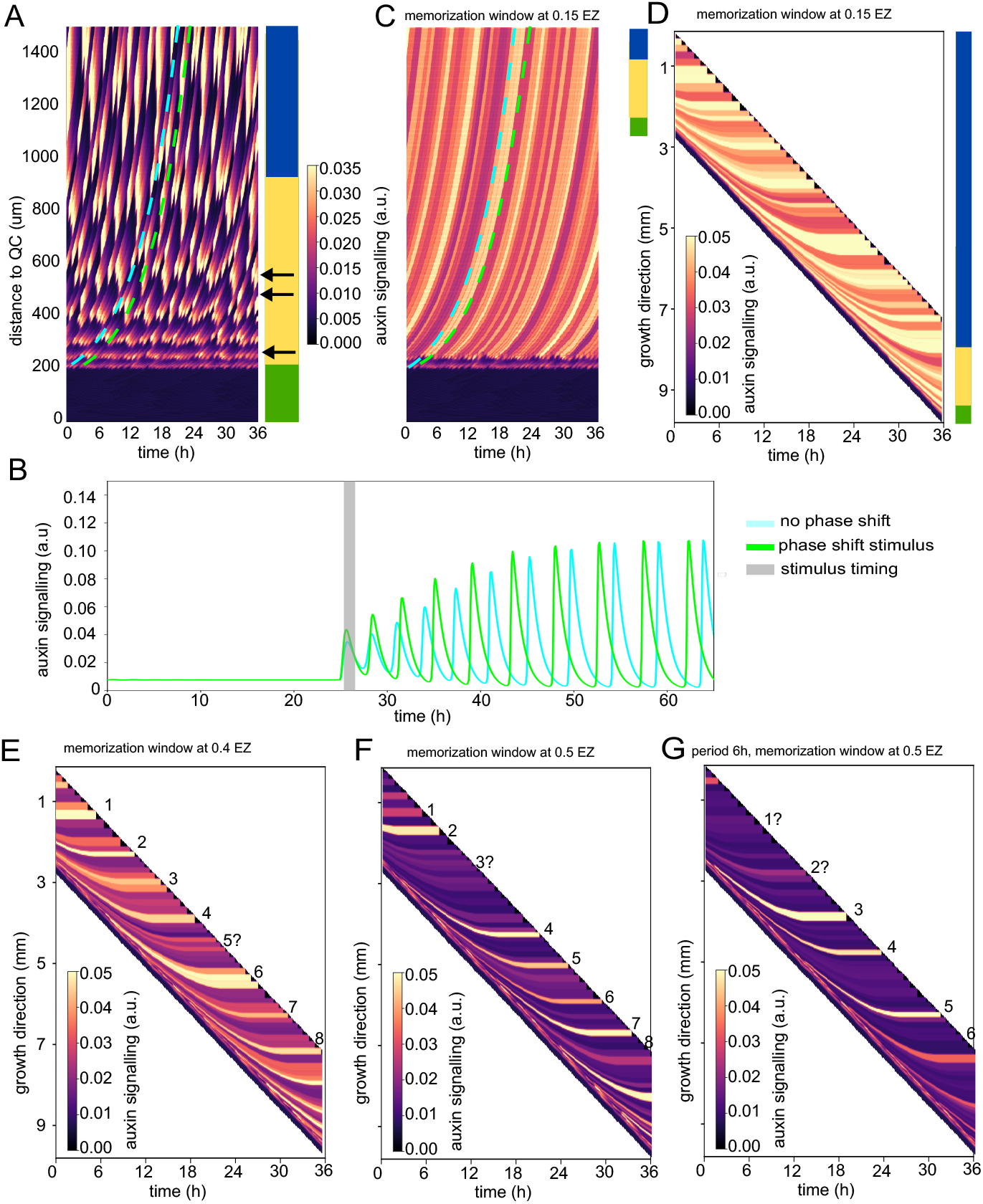
Periodic auxin input to EZ localised oscillations results in phase shifting. A) Kymograph depicting auxin signalling dynamics (ARF dimer level) for a situation where a 4.2h periodic increase of auxin is imposed in the early EZ (bottom 20%) and the parameter controlling the total amount of ARF in the system is increased from MZ to EZ such that oscillatory dynamics are only supported in the EZ and above. B) Temporal development of auxin signalling dynamics in a cell exposed to the auxin stimulus (cyan) and a cell not exposed to the auxin stimulus (green), with the timing of the auxin stimulus for the cyan cell indicated with the grey area. C) Kymograph depicting auxin signalling dynamics and subsequent memorisation from a position at 0.15 EZ (bottom arrow in A). D) Same kymograph as in B, but now for root tip displacing downward during growth. E) Downard oriented kymograph for situation where memorisation occurs from a position at 0.4 EZ (middle arrow in A). F) Downward oriented kymograph when memorisatin occurs from 0.5 EZ (top arrow in A). G) Downward oriented kymograph for a situation where auxin input is applied with a 6h instead of 4.2h period, with memorisation at 0.5 EZ. For E-G we indicated the hypothetical PBS, taken as elevations in memorized auxin signalling state with numbers.

Question marks are added if signalling level is less clearly elevated, likely resulting in unstable PBS.

Figure 10G shows auxin signalling and memorisation if an auxin stimulus period of 6h, slower than the inherent oscillator frequency, is used. The kymograph now shows 6 instead of 8 memorized high auxin peaks, indicating that stimulus frequency rather than endogeneous oscillator frequency dictates PBS formation rate. This begs the question what then the oscillator itself is for, although a role in translating signal modality or amplitude could be possible. Combined these results indicate that adding a periodic phase shifting signal to a EZ localized cell autonomous oscillator would result in a non-parsimonious, non-robust mechanism that furthermore requires multiple rounds of oscillations, inconsistent with experimental observations.

### Cryptic meristem oscillations are inconsistent with experimental data

A final seemingly logical hypothesis would be to propose the existence of oscillations in the meristem, which have not yet been picked up experimentally, which through time transform into the observed oscillations in the EZ. This would solve the problem of global time keeping and allow for differential phases in which cells arrive at the EZ. Indeed, for zebrafish somitogenesis oscilllations have been observed to increase in amplitude and decrease in frequency before their transformation into a temporally stable pattern (Shih et al., 2015). Thus, the question becomes what parameter change could explain the transition from unnoticed MZ to clear EZ oscillations. Note that this requires parameters that affect oscillation amplitude rapidly, given that only a single oscillation is observed in the EZ and before no oscillations are observed.

To investigate this I varied all 20 parameters of the simplified Middleton model between a low and high amplitude generating value . For the low amplitude value I take the lowest value still generating sustained oscillations. For the high amplitude value I take a value at or close to the default value described in Supplementary Table 2 for the oscillatory settings. For each parameter I recorded the mean level of auxin signalling (ARF dimer) for both the high and low amplitude value, the number of oscillation cycles needed to reach 95% of the new higher amplitude, the immediate and long term amplitude and the change in oscillation period (Supplementary Table 5).

For most parameters, amplitude increase requires a substantial number of oscillation cycles to fully unfold. Only for 3 parameters (*α*_*TIR*_, *p*_*m*_ and *d*_*auxin*_), oscillation amplitude overshot and then quickly equilibrated, allowing for a substantial oscillation amplitude increase for the first cycle. However, in all 3 cases, oscillation period decreased instead of increased, inconsistent with the prediction that oscillations should slow down before termination. Furthermore, for oscillations to not be observed in the MZ while being clearly observed in the EZ, at least 4 fold increase in oscillation amplitude should occur. Only 2 (*b*_*m*_ and *p*_*m*_ controlling AUX/IAA levels) out of the total of 20 parameters allowed for a 4 fold or higher oscillation amplitude increase, with one requiring multiple rounds of oscillations to unfold and the other resulting in oscillator speedup instead of slowing. Importantly, in both cases the transition from low to high amplitude oscillations involves a more than 2 fold increase in auxin signalling levels from the MZ to the EZ, inconsistent with experimental observations localizing the auxin signalling maximum in the MZ.

Of note, since parameters for low amplitude oscillations are close to the boundary of no longer supporting oscillations, combining two changes terminates oscillations rather than increasing amplitude differences. As a final attempt, I returned to the two ARFs two IAAs system, with the A system supporting and the B system not supporting oscillations. I changed the ratio between them from MZ to EZ from 1.35:0.25 A : B to 1.5:0.05 A : B to increase oscillation amplitude. Again, oscillation amplitude increase took 7 cycles to unfold and stayed below the threshold of a four fold increase.

## Discussion

Molecular clocks are an important method for time keeping in biological organisms, with the most well studied examples being the cell cycle and circadian clock present in single and multicellular organisms across the kingdoms of life. Another well studied example from developmental biology is the so-called somitogenesis clock, in which temporal oscillations in gene expression through growth become translated into a spatially periodic pattern of gene expression prepatterning future somite halves and boundaries. Hallmarks of clocks are their autonomous persistent oscillatory behavior in absence of external inputs, their potential for phase shifting and entrainment by external inputs and their potential to synchronize with other oscillators. Based on the observation of periodic gene expression variation, a root clock akin to the somitogenesis clock has been proposed to underly lateral root priming (Moreno et la., 2010). Still not all biological time keeping or sequentially ordered processes involve a clock, with a prominent example being the process of phyllotaxis, where auxin maxima prepatterning leaf primorodia are periodically formed from the interplay between tissue growth and auxin transport dynamics (Jönsson et al., 2006; Sassi & Vernoux, 2013; Smith et al., 2006).

Indeed, several differences between somitogenesis and lateral root priming beg the question of whether a cell-autonomous root clock is involved in the latter. Firstly, in the root cells seemingly undergo only a half “oscillation” -either an upstroke or downstroke-in contrast to the several rounds of oscillations that cells undergo in somitogenesis and that play an important role in somite patterning (McDaniel et al., 2024). Additionally, while in somitogenesis oscillations occur in the posterior growth region, in lateral root priming oscillations occur only once cells have left the meristem. It may in fact be more likely that lateral root priming and prebranch site formation resemble plant phyllotaxis rather than somitogenesis, with phyllotaxis also producing periodic auxin maxima at regular distances from the meristematic cells (Jönsson et al., 2006; Sassi & Vernoux, 2013; Smith et al., 2006). Note that also in phyllotaxis it would be expected that a constant distance from the SAM, periodic variations in gene expression will arise despite the absence of oscillations in single cells.

Other evolutionary considerations may also argue against the likelihood of a root clock. First, in animals, mobility is of critical importance for survival and therefore a robust, symmetric and proportional body plan is essential, explaining the evolution of a clock-work type mechanism. In contrast, in immobile plants flexible adjustment to environmental conditions rather than stereotypical patterning is of key importance to survive in a variety of different conditions. Thus, selective forces acting on animal body axis segmentation versus root system branching are likely very different (K. Ten Tusscher, 2020). Secondly, in animals with a segmented body plant the preferred solution appears to be a clock-and-wavefront type of segmentation, which likely has evolved in parallel not only in vertebrates but also in arthropods and annelids (Chipman, 2010; François et al., 2007; K. H. Ten Tusscher & Hogeweg, 2011). The simultaneous, hierarchical Drosophila segmentation is generally accepted as an evolutionary derived mode. In contrast, in plants expansion of the root system likely evolved from the bifurcating pattern observed in Lycophytes (Hetherington et al., 2020), and possibly via the merophyte based pattern in Ferns (Motte & Beeckman, 2019). Finally, in Cucurbites, lateral roots still arise in the meristem (Ilina et al., 2018) as in evolutionary older plant lineages, yet follow otherwise an Arabidopsis type route with founder cell specification. Thus, in plants clearly no repeatedly evolved and subsequently well conserved elongation zone oscillatory mechanism is present.

This current study provides yet another strong argument against the likelihood of a cell autonomous root clock underlying lateral root priming. First our study highlights the strong constraints on parameter conditions that apply for auxin signalling oscillations to occur, constraints that are further enhanced due to the presence of multiple AUX/IAA and ARF types typically being present in any plant tissue, as well as the presence of auxin exporters. Our study furthermore suggests that -assuming oscillations in gene expression and auxin (signalling) are occurring only in the (early) elongation zone-a clock mechanism is highly unlikely to underly lateral root priming. A critical difference between somitogenesis and lateral root priming identified here is the region in which oscillations occur. In the region where stem cell divisions and hence inheritance of cell states occur a global system level memory for oscillator phase can be established. In a zone without stem cell division, cells transition from a non-oscillatory to a oscillatory state from largely identical conditions, causing them to all start oscillations with the same phase and status and prohibiting variations in activity in the early elongation zone. Our study furthermore indicates that even with additional periodic input into the root clock specifically at the elongation zone, which would present a highly non-parsimonious mechanism, generating sufficient variation in early elongation zone activity is highly non-trivial. Finally, we show that the presence of a hidden oscillator in the meristem that only becomes apparent in the elongation zone is unlikely. In contrast, the reflux-and-growth mechanism we have proposed previously offers a parsimonious explanation for why oscillations occur specifically in the early elongation zone and in narrow vasculature cells (Van den Berg et al., 2021). Finally,on the level of entire root system architecture, we recently developed models simulating overall root system architecture (RSA) development to investigate the causes of differential root branching in brassinosteroid mutants (Khandal et al., 2025). In this study I demonstrated that growth based priming mechanisms such as reflux-and-growth or a Turing type mechanism could reproduce the reduced branching of brassinosteroid mutants. In contrast, a root clock mechanism, in which clock period is independent of root growth rate, would predict a highly branched architecture for brassinosteroid mutants, opposite to experimental observations.

I suggest that the previously reported periodic variations in gene expression (Moreno-Risueno et al., 2010) that have been interpreted as part of or downstream of a cell autonomous genetic oscillator are in fact all downstream of an emergent, non-cell autonomous process producing periodic variations in auxin signalling. Previously proposed candidate mechanisms are the periodic root cap apoptosis, reflected flow, as well as the reflux-and-growth mechanism (Mironova et al., 2010; van den Berg et al., 2021; Xuan et al., 2015a). Indeed, the fact that the periodic gene expression previously reported appears to consist of a sequence of waves containing different sets of genes appears more in line with a sequence of processes leading up to prebranch site formation set in motion by a periodic priming mechanism than that these gene expression waves are processes being part of the priming mechanism itself (Moreno-Risueno et al., 2010). Finally, I suggest that the negative feedback mechanism recently reported by Perianez-Rodriguez and co-workers is not a negative feedback giving rise to oscillations, but rather a negative feedback critical for auxin level homeostasis, which is necessary to produce regular spacing of prebranch sites. This is supported by their own modeling findings showing how in absence of this negative feedback an elevation of overall auxin signalling makes it hard to distinghuish peaks and valleys in auxin signalling and hence results in precocious prebranch site formation, and is consistent with the previously reported importance of an auxin signalling minimum in the lateral root forming region (Dubrovsky et al., 2011).

## Methods

### Single cell models

#### Original Middleton 2010 model

The original Middleton et al. 2010 model describes the dynamics of auxin, TIR receptors (TIR), auxin-TIR complexes (auxinTIR), Aux/IAA mRNA (M) and protein (A), auxin-TIR-AUX/IAA complexes (auxinTIRP), ARF monomers (A) and homodimers (A2), and ARF AUX/IAA dimers (AP), according to the equations given below. Note that we take the dedimensionalised equations, as derived in the paper. For an explanation of the meaning of parameters, values used, and units see Supplementary Table 1.

Given that the model assumes constant total levels of TIR (*α*_TIR_) and ARF (*α*_ARF_) it follows that:

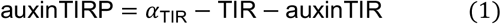

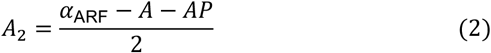

To describe the effect of ARF monomers (*F*_1_) and dimers (*F*_2_) on AUX/IAA gene expression dynamics the following functions are used:

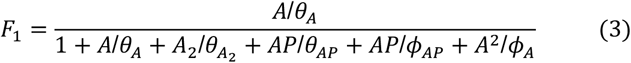

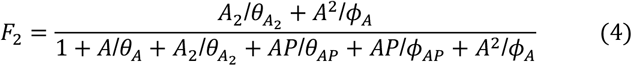

For the remaining variables the following equations are used:

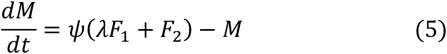

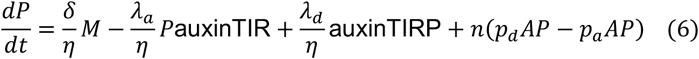

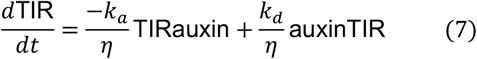

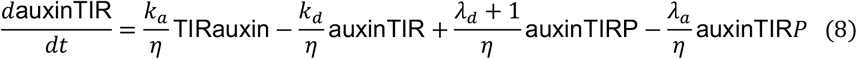

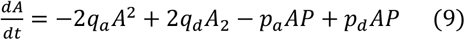

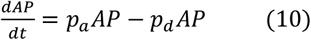

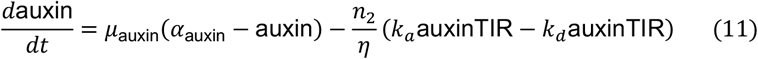

#### Simplified Middleton 2010 model

Conservation equation

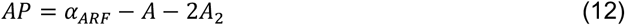

Note that I used the same conservation equation for total amount of ARF being constant, but instead of using it to write an algebraic expression for ARF homodimer levels I now use it to write an algebraic expression for ARF-AUX/IAA heterodimer levels. The reason for this different choice is that I wanted to have a differential equation of ARF homodimer levels and use their dynamics as a readout for auxin signalling levels in the models.

Helper functions (same as for full model):

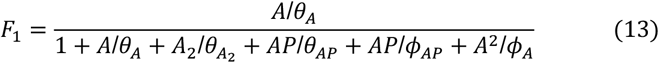

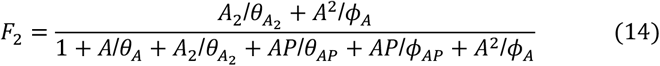

Differential equations:

To simplify the model and reduce the number of variables I made a number of assumptions. First, I assumed that changes in auxin level due to binding to TIR/AFB and subsequently AUX/IAA are small relative to changes from e.g. auxin transport and turnover and can thus be ignored. This enabled me to change auxin from a variable to a control parameter.

Next, I assumed that binding of auxin to TIR and of auxin-TIR complex to AUX/IAA protein is fast, enabling me to apply quasi-steady state approximations for equations 7 and 8 of the full model.

Solving 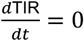 allows us to write:

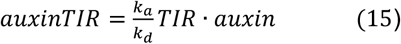

Solving 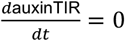 allows us to write

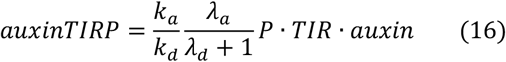

Substituting 15 into conservation equation 1 it follows that:

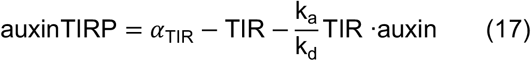

After reordering this gives us:

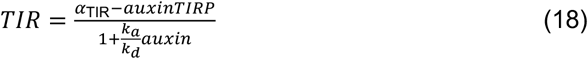

Substituting 18 back into 16 allows us to write:

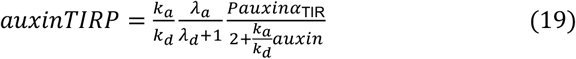

Finally from equation 11 of the full model it follows that the amount of AUX/IAA protein degraded through auxin-TIR binding is the 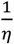 part of the 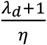 term, as it is the fraction of auxin-TIR-P complex that dissociates but does not return into the P equation.

Thus, decay of the AUX/IAA protein should be proportionate to the term I derived in equation 19. Finally, to prevent AUX/IAA protein levels to go to infinity in absence of auxin I added an auxin-independent baseline decay term.

Together these simplifications result in the following four variable model:

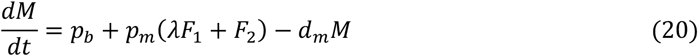

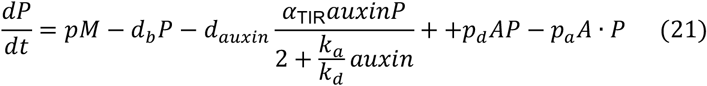

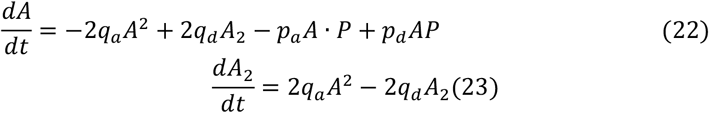

I find that for these equations, using the parameter settings that in the full model result in oscillations, no oscillations occurred. A bifurcation analysis along major parameters further confirmed that this simplified model could not support oscillations. I reasoned that by performing quasi steady state assumptions and making auxin TIR and AUX/IAA protein binding instantaneous some of the delays essential for oscillations were lost (Tyson and Novak, 2008). Next, I reasoned that in the current model, while auxin mediated degradation of AUX/IAA protein saturates with auxin also a saturation with protein level should occur as at some point TIR receptor levels become limiting. I therefore replaced equation 21 by the following equation:

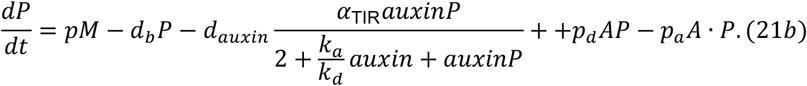

Which enabled oscillations in our simplified model for conditions similar to those in the full model. Note that a similar approach was followed in the Mellor et al., 2016 model (see below). The meaning and value of parameters can be found in Supplementary Table 2, for both oscillation supporting and non-oscillation supporting parameter settings used in the simulations.

#### Mellor 2016 model

I used the dedimensionalized version of the Mellor 2016 model. Note that the Mellor 2016 model does not consider ARF dimerization. In addition to the variables of the previous model, this model describes the dynamics of GH3 mRNA and protein (GH3m and GH3), LAX3 mRNA and protein (LAX3m and LAX3), mRNA and protein of an intermediate gene X (Xm and X), and of the auxin GH3 complex that forms prior to GH3 mediated auxin degradation (GH3aux).

Assuming a constant total level of ARF, in absence of ARF dimers the conservation equation in this model is:

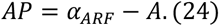

Helper functions:

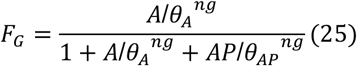

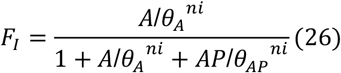

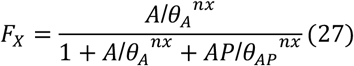

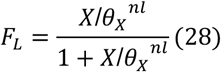

Differential equations:

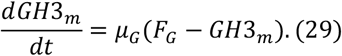

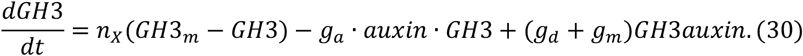

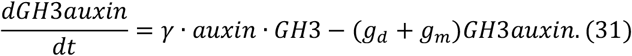

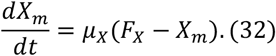

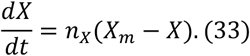

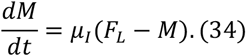

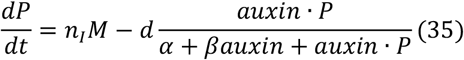

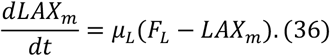

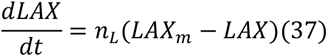

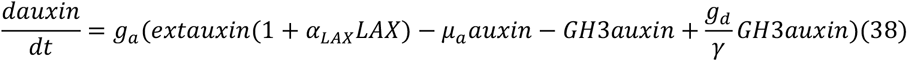

To study the effect of PIN mediated efflux the following equation was added:

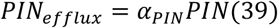

Where PIN level is either assumed constant or proportionate to X, changing equation 38 to:

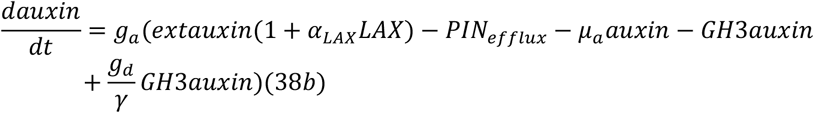

The meaning and value of parameter values can be found in Supplementary Table 3

### Dynamic 1D model

#### Layout & zonation

To appropriately describe protoxylem/pericycle dynamics withing a growing root tip I developed a 1D grid based tissue model consisting of a strand of cells. Each cell consists of a rectangle of grid points with a constant width, and a height that can change due to cytoplasmic growth, cell division and vacuolar expansion. Cells are positioned longitudinally and I superimpose at distinct positions the end of the proximal meristem (where cell division occurs), and the end of the transition zone or start of the elongation zone where slow cytoplasmic growth switches to rapid vacuolar expansion driven growth (Supplementary Table 4).

#### Growth division and elongation

Cell growth and expansion is modelled as in Mahonen et al. (2014). Briefly, if a cell undergoes either cytoplasmic growth or vacuaolar expansion a row of gridpoints is added to the shootward part of the cell and all shootward localized cels are shifted on row upward. Cell division occurs if cells have reached double their initial size.

#### Growth parametrisation

I parametrised initial cell height, cytoplasmic growth rates and hence cell size doubling and division time, proximal meristem size and the boundary between the transition and elongation zone such that I could approximate the root growth dynamics described in Goh et al. (2023) (Supplementary Table 4). I simulated a total of 2654 μm of tissue, encompassing meristem, elongation zone, and a substantial part of the early differentiation zone.

Each cell has a differentiation variable, described by a differential equation. Initially the value of this variable is zero, and it only starts increasing once cells enter the elongation zone. Parameters of the differentiation variable are tuned such that the differentiation variable exceeds a threshold level of 85 at around 7hrs, beyond which expansion ceases and cells no longer grow:

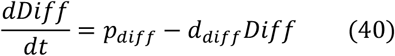

See Supplementary Table 4 for parameter values.

To incorporate some stochasticity, instead of making all cells divide synchronously at a constant rate, I incorporated a bottom stem cell and initial cell, with slower division rates than the rest of the meristematic transit amplifying cells (Supplementary Table 4). Additionally, each time a cell divides, daughter cells receive a division time in between 95% and 105% of the average division time for their respective position.

#### Intracellular dynamics & parametrisation

Each cell contains the auxin signalling network as described by the simplified Middleton 2010 model. All cells are parametrized identically using the values described in Supplementary Table 2, except for the total amount of ARF, which is used to control the capacity of the cells to oscillate. In our simulations I use either high ARF, oscillation promoting, levels (1.5) in the meristem and transition zone or above the transition zone. Elsewhere low ARF, non-oscillation promoting levels (0.5) are used. Upon cell division, daughter cells inherit the state of their mother cell.

#### Special settings

Superimposed mRNA oscillations (Fig 10):

For these simulations we replaced equation 21 with:

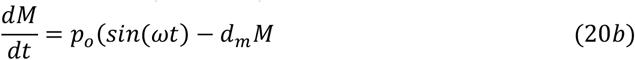

With *p*_*o*_ = 0.125*p*_*m*_ to compensate for the fact that a sine function has a maximum of 1 while *λF*_1_ + *F*_2_ typically stays below 0.35, and *ω* = 1.566 *radians*/*h*. From *ω* = 2*πf* it then follows that *f* = 0.25 *cycles*/*h* resulting in a period of 4 hours.

Periodic stimuli (Fig 11):

For these simulations we replace the constant value of auxin by the equation:

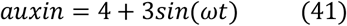

where the values 4 and 3 were chosen to have auxin input vary between 1 and 7, and using for *ω* either a value of 1.50 or 1.05 *radians/h*, and hence f a value of 0.238 or 0.167 *cycles/h*, resulting in a period of 4.2h or 6h

#### Finite domain size

Cells exceeding a certain threshold distance from the start of the tissue are removed from the simulation. This enables us to maintain a constant simulation domain size despite continuous growth.

#### Numerics

ODEs are solved using simple Euler forward integration with a timestep of 0.025s. Spatial resolution of the grid was 2 microm.

### Model simulation and analysis

The single cell full Middleton 2010 model, simplified Middleton 2010 model and Mellor 2016 model were coded in python, and numerically solved using the function odeint (Virtanen et al., 2020). These codes were used to generate the single cell temporal dynamics shown in Figures 4-6. Additionally, for bifurcation analysis (Fig 4) the full and simplified Middleton models were coded in R, making use of the package custom script Grind.R (created by Rob J. de Boer, for information and download see https://bioinformatics.bio.uu.nl/rdb/grind.html), which in turn uses the package deSolve for numerically solving the equations (Soetaert et al., 2010).

The 1D model was coded in C++, using simple Euler forward integration with a timestep of 0.25s and a space step of 1μm. The spatiotemporal data generated by this code was plotted using separate python codes.

## Author contributions

K.t.T conceived the study, constructed the models, ran simulations and analysed results and wrote the manuscript.

## Acknowledgements

I am indebted to Bas van den Herik and Benjamin Planterose Jiminez for discussions, feedback and technical support.

## Data and resource availability

All model codes used in this study and an explanatory Readme file will be made available upon publication on the groups github page: https://github.com/kirstentt

Shown are the parameters of the simplified Middleton model developed in this study that were varied to obtain oscillation amplitude differences. Note that *θ*_*A*_ is left out from the table because of its limited effect on oscillation amplitude.

## Supplementary Figures

**Supplementary Figure 1.**
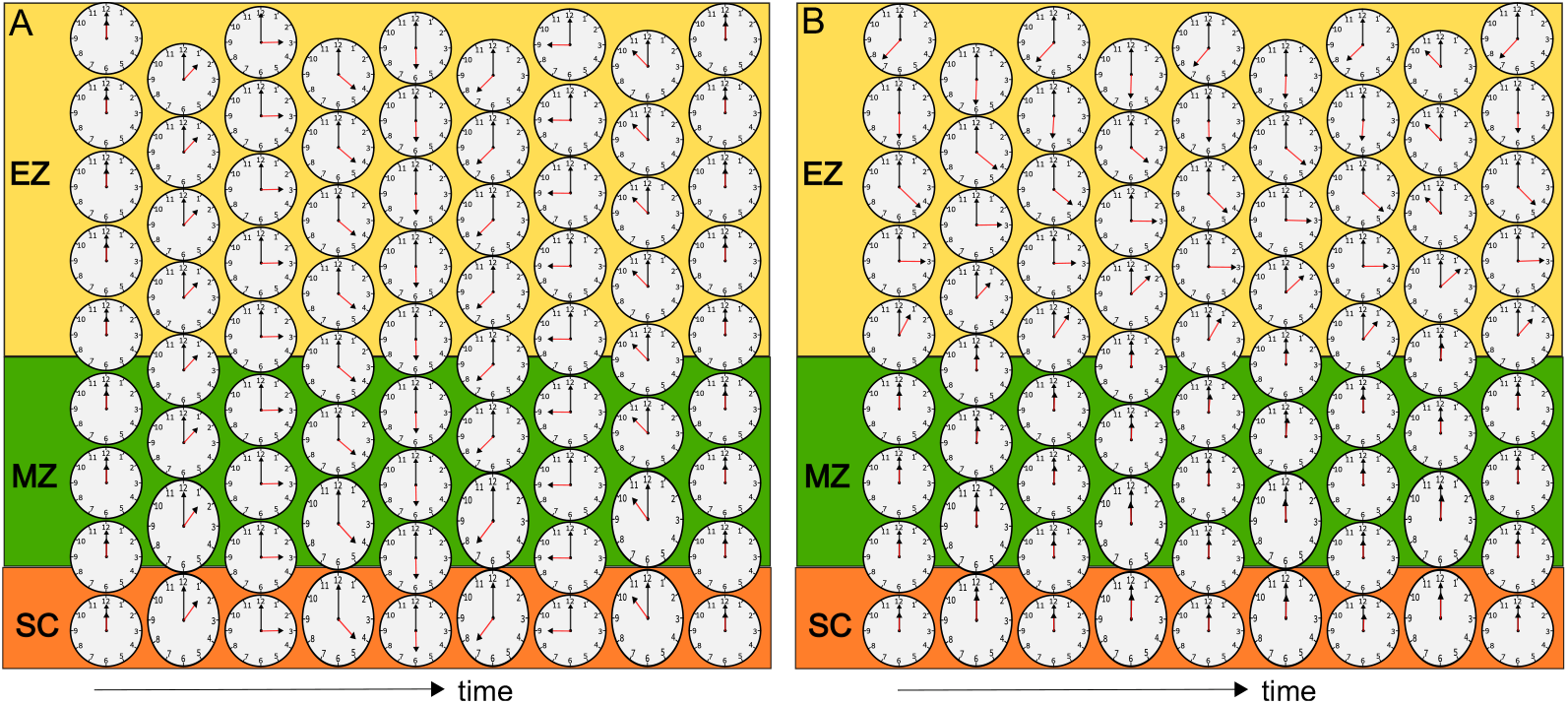
Clockwork versus stopwatch like oscillatory behavior. A) If oscillations occur in the stem cell niche and meristem, through inheritance of oscillator phase after division (see growing and dividing clocks) a global time keeping arises (look at clock state in bottom row from left to right), and cells arriving at different times do so with different clock phases. B) If oscillations start in the elongation zone, all cells in the stem cell niche and meristem are in a steady state, like a stopwatch not yet pressed to start, and all cells start with the same clock state when entering the elongation zone, like when crossing the start and a stopwatch is pressed to run.

**Supplementary Table 1.** Parameters for the original Middleton 2010 model. Parameter values of the dedimensionalised model as derived in Middleton et al. (2010) are used, hence parameters are dimensionless. Symbols, meaning and values are provided.

| *Parameter* | *Meaning* | *Value* |
| --- | --- | --- |
| $\alpha_{TIR}$ | Total TIR amount | 1.0 |
| $\alpha_{ARF}$ | Total ARF amount | 1.0 |
| $\theta_A$ | Affinity constant for A | 0.1 |
| $\theta_{A2}$ | Affinity constant for A2 | 0.01 |
| $\theta_{AP}$ | Affinity constant for AP | 0.1 |
| $\varphi_{AP}$ | Cooperativity constant for AP | 0.1 |
| $\varphi_A$ | Cooperativity constant for A | 0.1 |
| $\psi$ | mRNA production scaling | 100 |
| $\lambda$ | Weight of F1 in M dynamics | 0.1 |
| $\delta$ | Protein production rate | 1 |
| $\eta$ | Scaling factor for protein/complex dynamics | 0.01 |
| $\lambda_a$ | AuxinTIR P association constant | 10 |
| $\lambda_d$ | AuxinTIRP dissociation constant | 1 |
| $n$ | Stoichiometric coefficient | 1 |
| $p_a$ | AP association constant | 100 |
| $p_d$ | AP dissociation constant | 100 |
| $k_a$ | Auxin–TIR association constant | 0.1 |
| $k_d$ | Auxin–TIR dissociation constant | 1 |
| $q_a$ | ARF dimerization rate | 1 |
| $q_d$ | ARF dimer dissociation rate | 1 |
| $\mu_{auxin}$ | Auxin turnover rate | 10 |
| $\alpha_{auxin}$ | Relative auxin production constant | 1 |
| $n_2$ | Stoichiometric factor in auxin–TIR dynamics | 10 |

**Supplementary Table 2.**
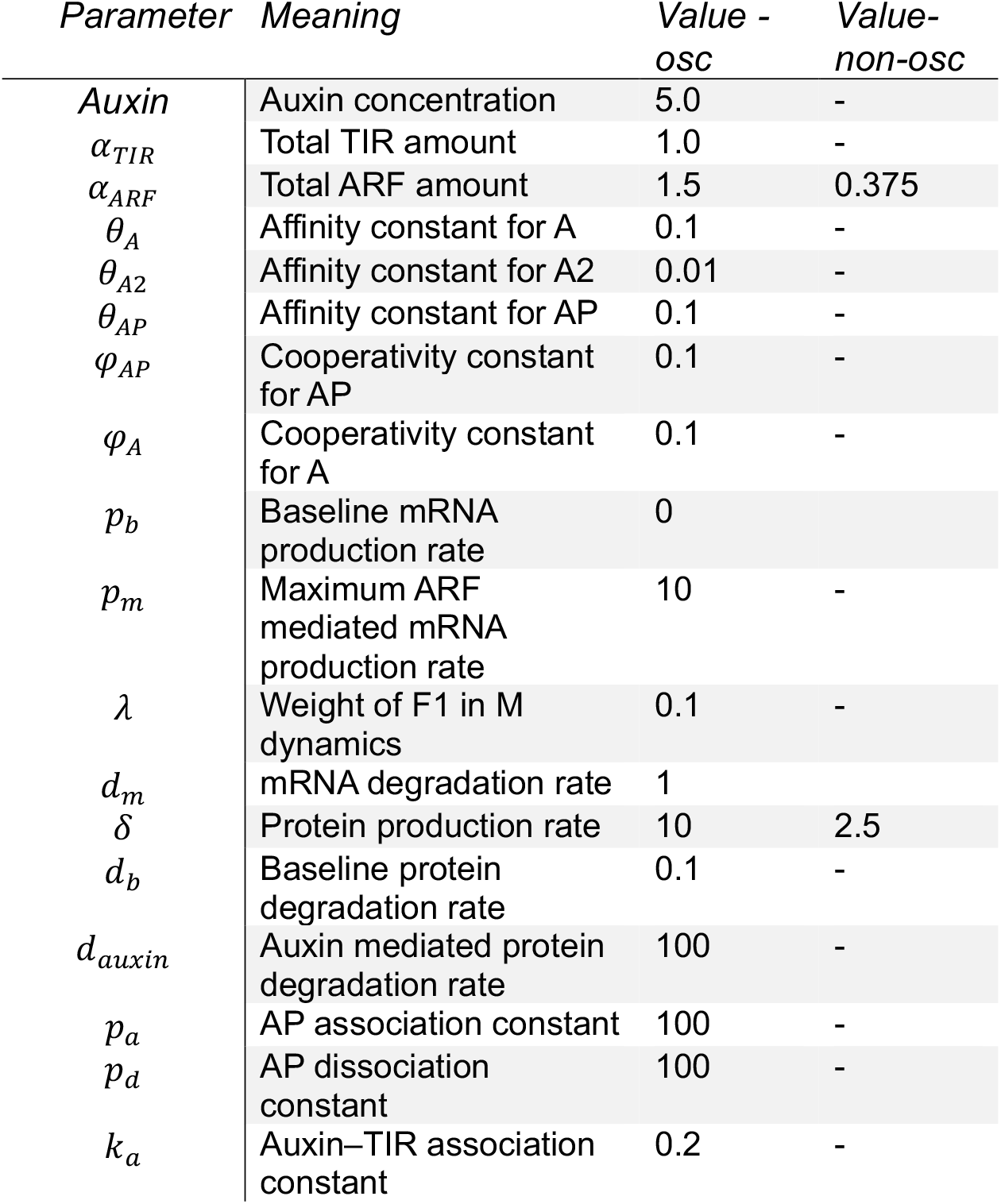
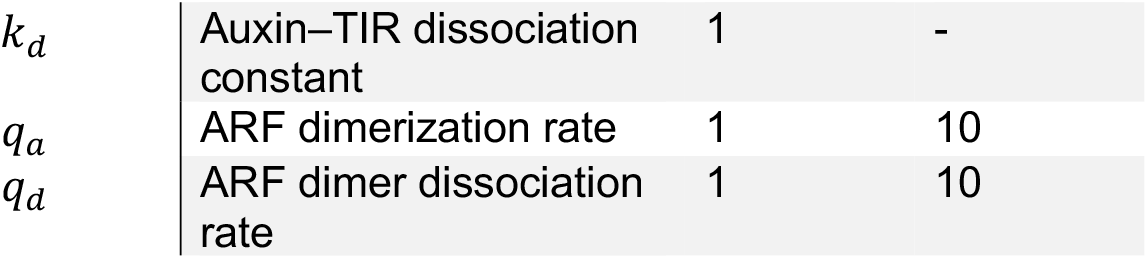
parameters for the simplified Middleton 2010 model. “Value-osc” are the parameter values used to simulate oscillatory dynamics. “Value-non-osc” are parameter values to generate non-oscillatory dynamics. “-” indicates parameter values identical to the oscillatory regime are used, only distinct parameter values are provided for clarity.

**Supplementary Table 3.**
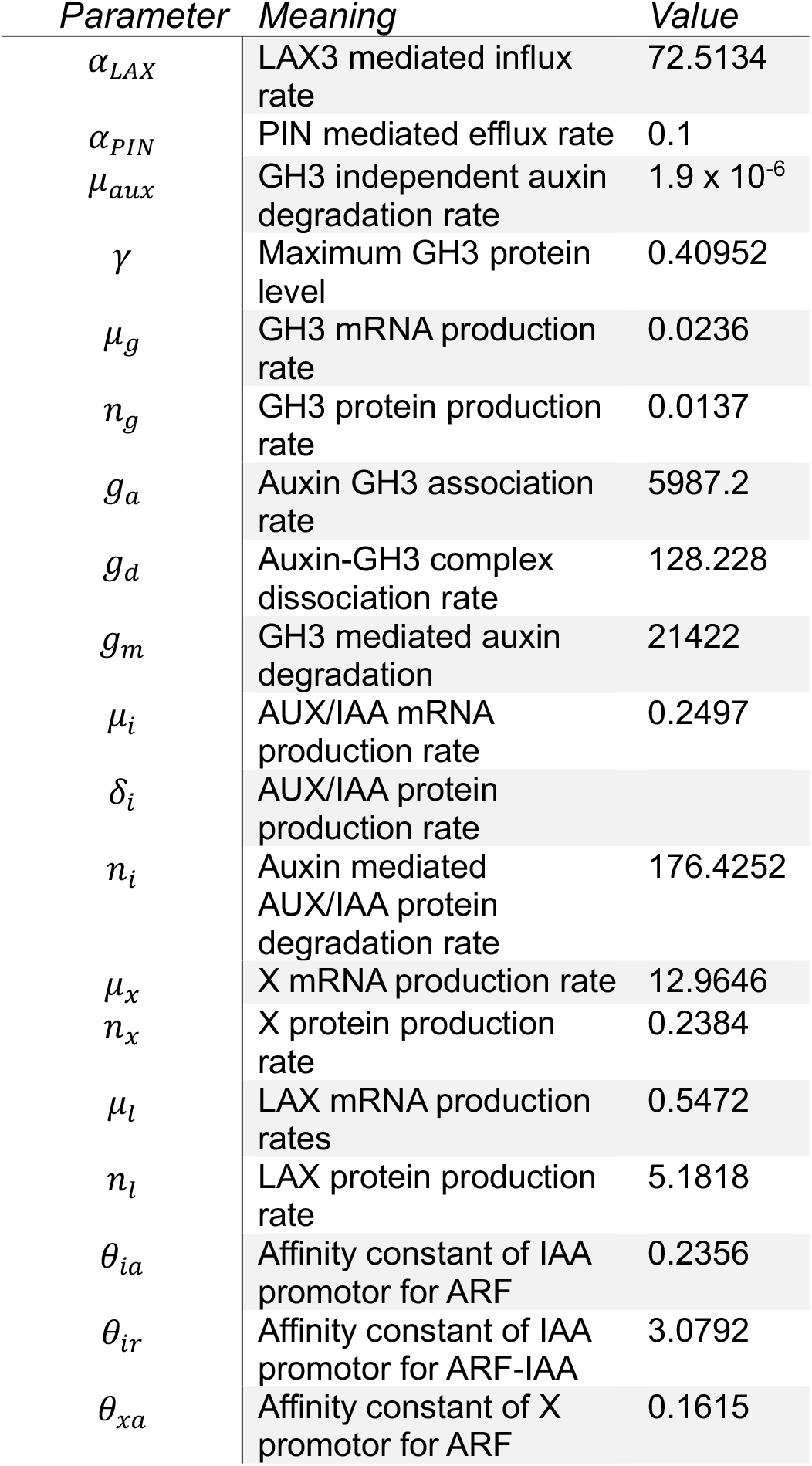
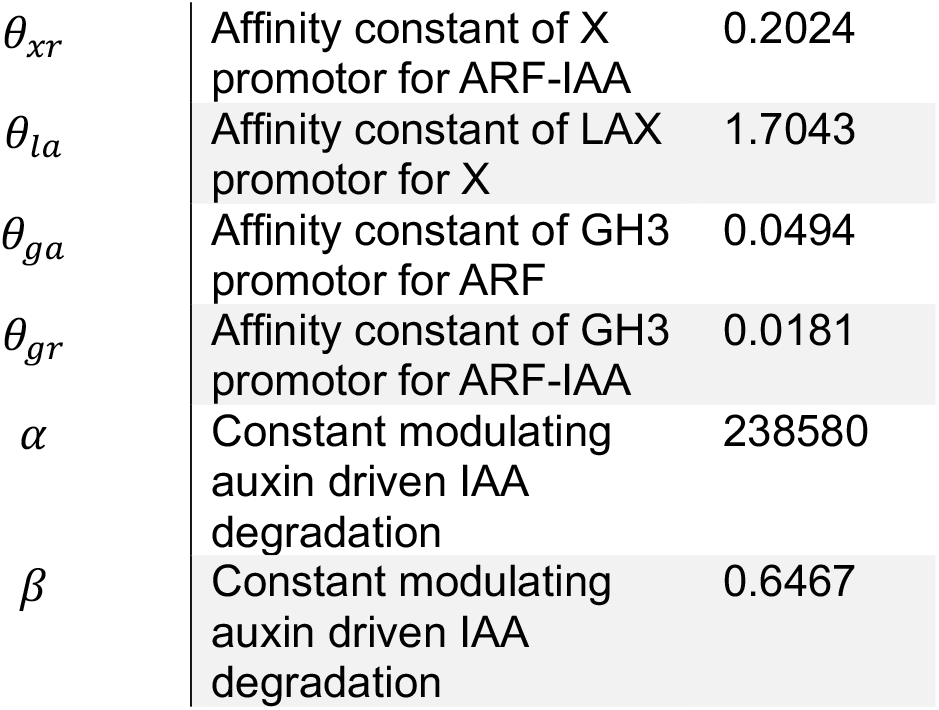
Parameters for the Mellor 2016 model. Note that as for the original and simplified Middleton model, the dedimensionalized version of the model is used, resulting in dimensionless parameters. As before symbols, meaning and values are provided.

| *Parameter* | *Meaning* | *Value* |
| --- | --- | --- |
| $\alpha_{LAX}$ | LAX3 mediated influx rate | 72.5134 |
| $\alpha_{PIN}$ | PIN mediated efflux rate | 0.1 |
| $\mu_{aux}$ | GH3 independent auxin degradation rate | $1.9 \times 10^{-6}$ |
| $\gamma$ | Maximum GH3 protein level | 0.40952 |
| $\mu_g$ | GH3 mRNA production rate | 0.0236 |
| $n_g$ | GH3 protein production rate | 0.0137 |
| $g_a$ | Auxin GH3 association rate | 5987.2 |
| $g_d$ | Auxin-GH3 complex dissociation rate | 128.228 |
| $g_m$ | GH3 mediated auxin degradation | 21422 |
| $\mu_i$ | AUX/IAA mRNA production rate | 0.2497 |
| $\delta_i$ | AUX/IAA protein production rate | |
| $n_i$ | Auxin mediated AUX/IAA protein degradation rate | 176.4252 |
| $\mu_x$ | X mRNA production rate | 12.9646 |
| $n_x$ | X protein production rate | 0.2384 |
| $\mu_l$ | LAX mRNA production rates | 0.5472 |
| $n_l$ | LAX protein production rate | 5.1818 |
| $\theta_{ia}$ | Affinity constant of IAA promotor for ARF | 0.2356 |
| $\theta_{ir}$ | Affinity constant of IAA promotor for ARF-IAA | 3.0792 |
| $\theta_{xa}$ | Affinity constant of X promotor for ARF | 0.1615 |
| $\theta_{xr}$ | Affinity constant of X<br>promotor for ARF-IAA | 0.2024 |
| $\theta_{la}$ | Affinity constant of LAX<br>promotor for X | 1.7043 |
| $\theta_{ga}$ | Affinity constant of GH3<br>promotor for ARF | 0.0494 |
| $\theta_{gr}$ | Affinity constant of GH3<br>promotor for ARF-IAA | 0.0181 |
| $\alpha$ | Constant modulating<br>auxin driven IAA<br>degradation | 238580 |
| $\beta$ | Constant modulating<br>auxin driven IAA<br>degradation | 0.6467 |

**Supplementary Table 4.** Parameters for the growing1D model.

| *Parameter* | *Meaning* | *Value* | *Units* |
| --- | --- | --- | --- |
| $B_{MZ-TZ}$ | Boundary between<br>proximal meristem and<br>transition zone | 155 | microm |
| $B_{TZ-EZ}$ | Boundary between<br>transition zone and<br>elongation zone | 200 | microm |
| $H_{init}$ | Initial cell height | 4 | microm |
| $H_{div}$ | Cell height at which<br>division takes place | 9 | microm |
| $H_{max}$ | Maximum cell height<br>used to compute<br>elongation rate | 100 | microm |
| $t_{div}$ | Time it takes for transit<br>amplifying cell to grow<br>from $H_{init}$ to $H_{div}$ and<br>undergo division | 11 | h |
| $t_{stem}$ | Division time of stem<br>cell | 30 | h |
| $t_{initial}$ | Division time of initial<br>cell | 17 | h |
| $t_{elong}$ | Time it takes for a cell to<br>grow from $2H_{init}$ to<br>$H_{max}$ | 18 | h |
| $T_{diff}$ | Differentiation threshold<br>beyond which cells no<br>longer elongate | 85 | a.u. |
| $p_{diff}$ | Rate of increase of<br>differentiation variable | 0/0.0024 | s-1 |
| $d_{diff}$ | Turnover rate of<br>differentiation variable | 0.000024 | s-1 |

**Supplementary Table 5.**
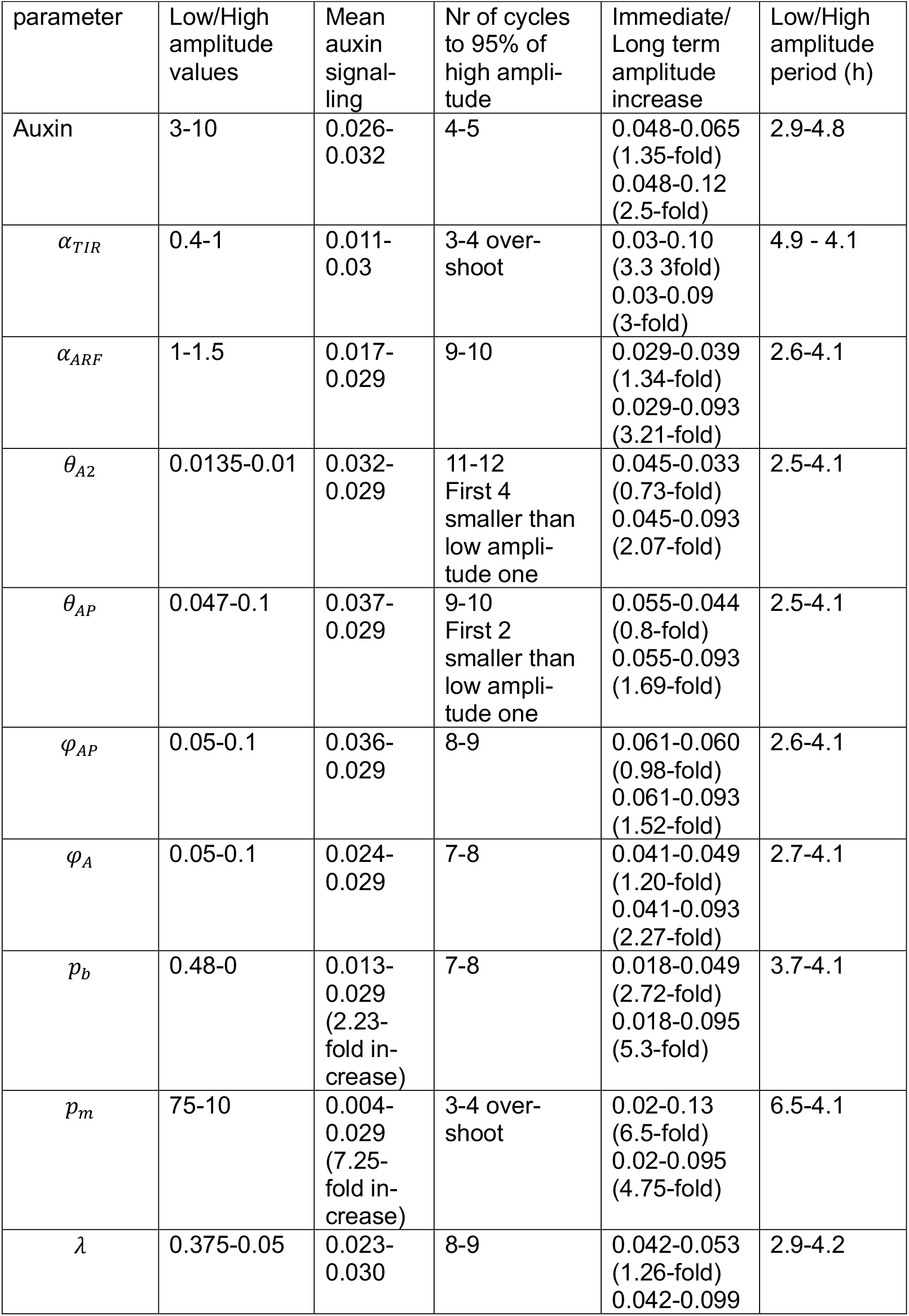
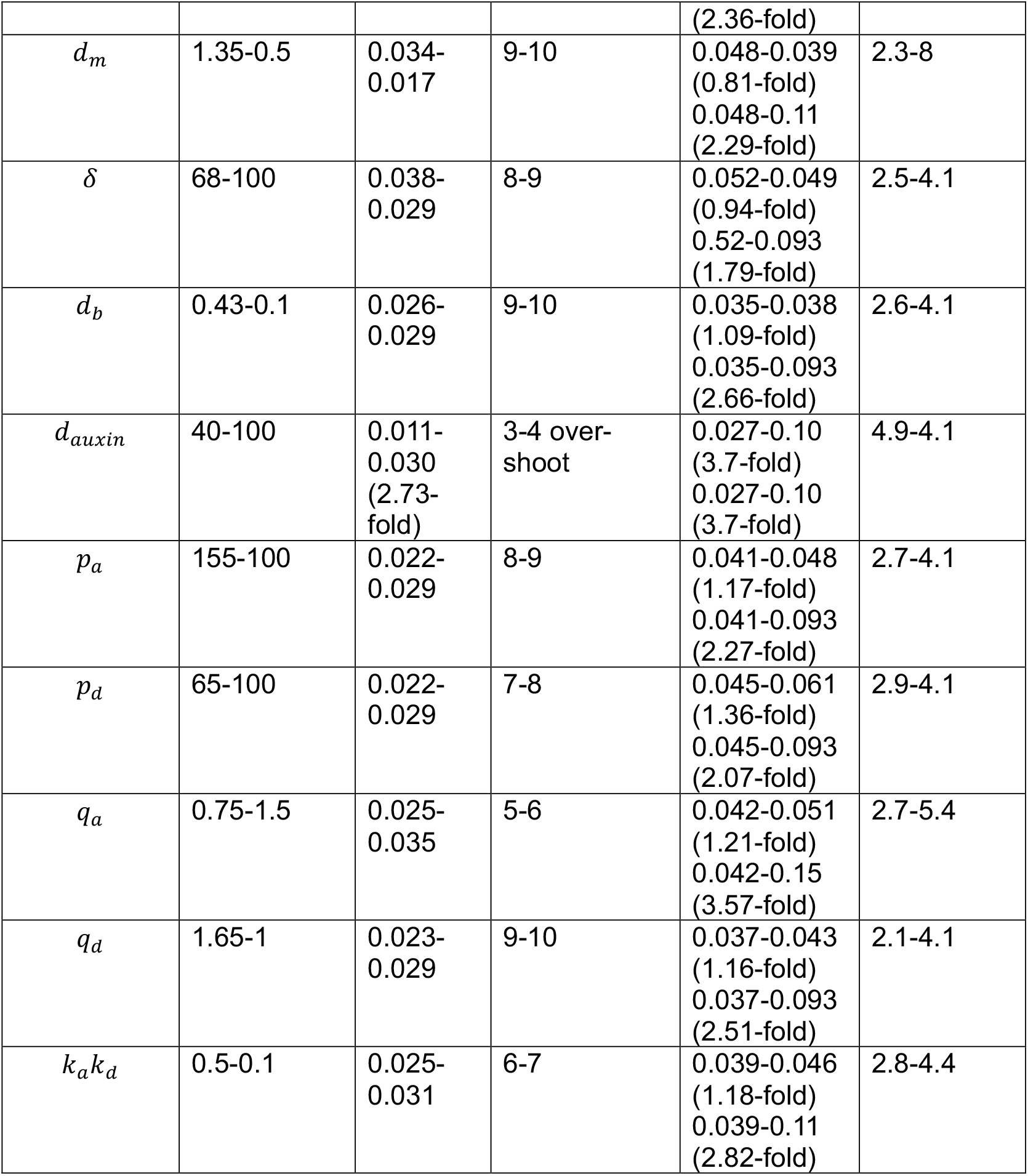
Effect of parameters on oscillation characteristics.

